# The Deformability of the Mammalian Cell Nucleus is Determined by the Identity of the Lamin Rod Domain

**DOI:** 10.1101/2025.07.22.666133

**Authors:** Jacob Odell, Yuliang Tang, Yogeshwari Sanjayrao Ambekar, Gururaj Rao Kidiyoor, Hassan Saadi, Graeme F. Woodworth, Liam J. Holt, Giuliano Scarcelli, Haiyuan Yu, Jan Lammerding

## Abstract

Lamins are nuclear intermediate filament proteins with diverse functions, ranging from organizing chromatin and regulating gene expression to providing structural support to the nucleus. Mammalian cells express two types of lamins, A-type and B-type, which, despite their similar structure and biochemical properties, exhibit distinct differences in expression, interaction partners, and function. One major difference is that A-type lamins have a significantly larger effect on the mechanical properties of the nucleus, which are crucial for protecting the nucleus from cytoskeletal forces, enabling cell migration through confined spaces, and contributing to cellular mechanotransduction. The molecular mechanism underlying this difference has remained unresolved. Here, we applied custom-developed biophysical and proteomic assays to lamin-deficient cell lines engineered to express specific full-length lamin proteins, lamin truncations, or chimeras combining domains from A- and B-type lamins, to systematically determine their contributions to nuclear mechanics. We found that although all expressed lamins contribute to the biophysical properties of the nuclear interior and confer some mechanical stability to the nuclear envelope, which is sufficient to protect the nuclear envelope from small cell-intrinsic forces and ensure proper positioning of nuclear pores, A-type lamins endow cells with a unique ability to resist large forces on the nucleus. Surprisingly, this effect was conferred through the A-type lamin rod domain, rather than the head or tail domains, which diverge more substantially between A- and B-type lamins and play important roles in lamin network formation. Collectively, our work provides an improved understanding of the distinct functions of different lamins in mammalian cells and may also explain why mutations in the A-type lamin rod domain cause more severe muscle defects in mouse models than other mutations.

## Introduction

Lamins are nuclear intermediate filament proteins that are the principal components of the nuclear lamina, a dense protein meshwork at the inner surface of the inner nuclear membrane (INM). Like other intermediate filaments, lamins have a conserved tripartite domain organization consisting of a small N-terminal head domain, a central helical rod domain, and a C-terminal tail that contains a nuclear localization signal and an Immunoglobulin-like (Ig-like) fold^1,2^. Lamin monomers dimerize via their rod domains to form a coiled-coil, and dimers form anti-parallel tetramers that assemble into protofilaments via head-to-tail polymerization of the coiled-coil domains, with lateral association of protofilaments resulting in the final filaments^3^. While the rod domain is essential for dimerization via coiled-coil formation and necessary and sufficient for the assembly of filaments, the head and tail domains are thought to contribute to longitudinal head-to-tail assembly of lamin dimers^4–6^. Vertebrate lamins are grouped into A-type and B-type lamins, which differ in their expression patterns, biochemical properties, and posttranslational modifications^7^. The major A-type lamins in mammalian cells are lamins A and C (LaA and LaC, respectively), which are alternatively spliced products of the *LMNA* gene. The major B-type lamins are lamin B1 (LaB1), encoded by *LMNB1*, and lamin B2 (LaB2), encoded by *LMNB2*. The different lamin types likely evolved as a result of paralog duplication concomitant with two rounds of whole genome duplication in the vertebrate lineage, which allowed for the A-type and B-type lamins to diverge from one another in terms of both sequence and function^8–10^.

Mammalian lamin types share conserved amino acid residues in their primary sequence (48% identity, 66% similarity between human LaA and LaB1) (Extended Data Figure 1), and the different lamin types have several functions in common. Both A-type and B-type lamins localize to the nuclear periphery and interact with chromatin and transcriptional regulators to influence genome organization and gene expression^11^. Nonetheless, A-type and B-type lamins, which form separate filament systems^12,13^, exhibit important differences in expression and function. A- and B-type lamins are differentially expressed during development^14^, with B-type lamins being ubiquitously expressed across tissues, whereas A-type lamins are highly enriched in stiff and mechanically loaded tissues, such as skeletal muscle and the heart^15^. Nearly all human diseases that involve mutations or abnormal expression of lamins (collectively termed laminopathies) involve alterations in A-type rather than B-type lamins^16^. A-type lamins are generally considered to be the major contributor to nuclear stiffness, because depletion of LaA and LaC leads to nuclei that are more deformable and mechanically more fragile than nuclei of wild-type cells^17–28^. The role of A-type lamins in determining nuclear deformability and stability is not only crucial in the pathogenesis of laminopathies and maintaining nuclear integrity in muscle tissues, but also in the ability of cells to migrate through tight interstitial spaces during development and cancer metastasis, and in the ability of cells to respond to mechanical stimuli^29–32^.

On the other hand, loss of Lamin B1 has a smaller effect on nuclear deformability^19,33^, yet loss of Lamin B1 also compromises nuclear envelope integrity and leads to increased rates of nuclear blebbing and nuclear envelope rupture^19,22,34–37^. It remains unclear which domains of the A-type and B-type lamins determine their discrete contributions to the mechanical properties of the nucleus.

To address this question, we designed a complementation system to exogenously express full-length, truncated, or chimeric lamins in lamin-deficient cells and determine the effect of each lamin type on nuclear deformability, as well as the biophysical and biochemical factors that modulate nuclear mechanics. We found that all lamins contribute to the mechanical stability of the nuclear envelope and modulate the biophysical properties of the nuclear interior. While this contribution is sufficient to protect the nuclear envelope from small, cell-intrinsic forces, A-type lamins equip cells with the ability to better resist large forces on the nucleus. We identified that this unique ability of A-type lamins is primarily conferred through their rod domain, rather than their head or tail domains. These results implicate the emergent lamina network formed by lamins containing the A-type rod domain as a key driver for the difference between lamin types in providing structural support to the nucleus, which is crucial to resisting cytoskeletal forces in muscle and other mechanically stressed cells and tissues.

## Results

### Micropipette aspiration reveals different stiffness conferred by A-type and B-type lamins

To evaluate the contributions of different lamins and domains to nuclear mechanical stability, we used a recently developed high-throughput micropipette aspiration system^38^ that quantifies nuclear stiffness by measuring the deformation of the nucleus into a small aspiration channel as cells in suspension are subjected to an external pressure gradient inside a microfluidic device (Figure 1A). Micropipette aspiration is ideally suited to evaluate the contribution of lamins to nuclear mechanics because it imposes large deformations on the nucleus, which are primarily resisted by the nuclear lamina^23^. Furthermore, the use of live, intact cells avoids potential damage to the nuclear envelope during nuclear isolation and altering nuclear stiffness due to changes in multi-valent ion concentrations^39,40^. We first applied this technique to mouse embryo fibroblast (MEFs) lacking either lamin A/C (*Lmna*^-/-^), lamin B1 ((*Lmnb1*^-/-^), or ‘triple lamin knockout’ (TKO, i.e., *Lmna^-/-^ Lmnb1^-/-^ Lmnb2^-/-^*) MEFs, along with wild-type controls. These cell lines have been well characterized previously for their nuclear mechanical properties^17–19,36^ and represent an excellent model to study the contributions of individual lamin proteins to cell biology. Since previous studies had shown very similar effects of lamin B1 and B2 depletion on nuclear mechanics^41^, we did not assess *Lmnb2^-/-^* MEFs in our studies.

**Figure 1:**
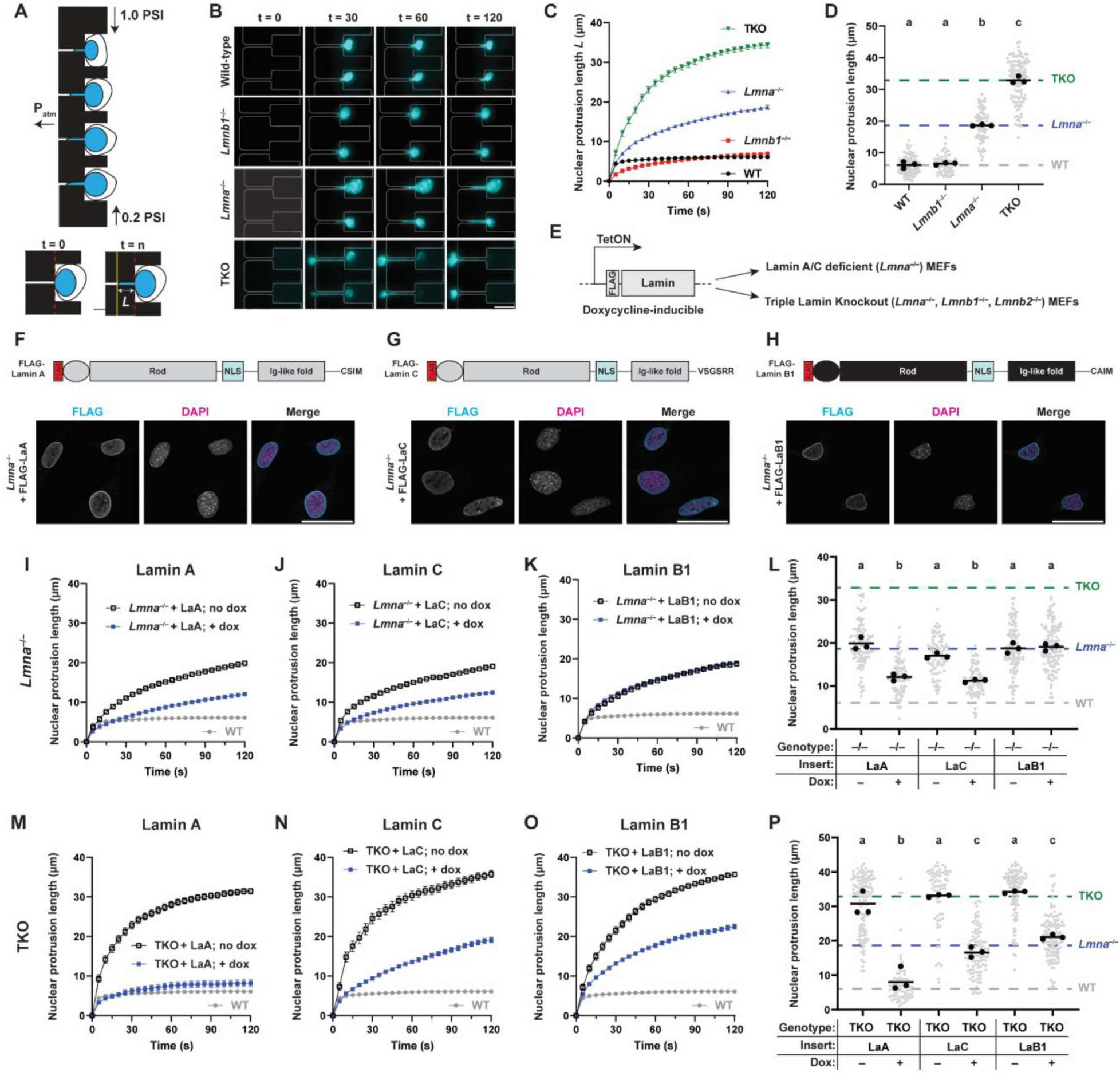
Contributions of A-type and B-type lamins to nuclear mechanics revealed by micropipette aspiration. (**A**) Schematic of microfluidic micropipette aspiration device. Nuclear stiffness in intact cells is inferred from the deformation of the nucleus into the small aspiration channel under an applied pressure gradient. (**B**) Images from representative time series of cells subjected to micropipette aspiration. Nuclei of cells of the indicated genotypes were stained with Hoechst 33342 and aspirated into the device under defined pressure. Scale bar = 20 µm. (**C**) Quantification of nuclear protrusion over time of the cells shown in (B). Points represent the average protrusion of nuclei at a given time point, error bars represent s.e.m. (**D**) Nuclear protrusion lengths measured 100 s after the start of aspiration. Grey points indicate measurements from individual cells, black points indicate replicate means, and bars indicate overall means. Sets of points with the same letter above them are not significantly different, whereas different letters indicate *p* < 0.05, One-way ANOVA with Tukey’s multiple comparison test. (**E**) Exogenous expression strategy to introduce a lamin construct of interest under the control of a doxycycline-inducible promoter. Different vectors are used to express either LaA, LaC, or LaB1, or, in later experiments, lamin truncations and chimeras, in either lamin A/C-deficient (*Lmna*^-/-^) or triple lamin knockout (TKO) MEFs. (**F-H**) Schematic representation of LaA, LaC, or LaB1 constructs, showing the different lamin domains (top), and representative immunofluorescence images of *Lmna*^-/-^ MEFs expressing each lamin protein (bottom). Scale bar = 50 µm. (**I-K**) Quantification of nuclear protrusion over time for *Lmna*^-/-^ MEFs expressing LaA (I), LaC (J), or LaB1 (K), as in (**C**). Grey points from the wild-type control are included for reference on each plot comparing the baseline deformability (no dox), to the deformability of cells expressing the exogenous lamin. (**L**) Quantification of *Lmna*^-/-^ MEFs expressing full-length lamins 100 s after the start of aspiration, as in (D). (**M-O**) Quantification of nuclear protrusion over time for TKO MEFs expressing LaA (M), LaC (N), or LaB1 (O). (**P**) Quantification of TKO MEFs expressing full-length lamins 100 s after the start of aspiration, as in (D).

Consistent with previous reports^28,42^, *Lmna*^-/-^ MEFs had more deformable nuclei than wild-type cells, evidenced by increased nuclear protrusion into the aspiration channel over time (Figure 1B-D). In contrast, loss of LaB1 did not significantly alter nuclear deformability compared to wild-type controls. TKO MEFs had even more deformable nuclei than *Lmna^-/-^* MEFs, with several TKO cells passing completely through the aspiration channel during the observation period (Figure 1B). These findings indicate that although loss of LaB1 does not impair nuclear stability in the presence of A-type lamins, B-type lamins contribute to nuclear stability in the absence of A-type lamins.

To further characterize the contributions of individual lamin proteins to nuclear mechanical stability, we developed an exogenous expression system to express specific FLAG-tagged lamins at consistent levels in either *Lmna*^-/-^ or TKO MEFs (Figure 1E) and assess the extent to which each lamin rescued nuclear deformability. Expression of the lamin constructs is driven by a doxycycline (dox)-inducible promoter so that the protein is expressed only following addition of doxycycline, allowing for control over the timing and levels of expression (Extended Data Figure 2) and enabling comparison to the baseline nuclear deformability of each cell line (‘no dox’ condition). Immunofluorescence staining for the FLAG tag attached to the exogenously expressed lamins confirmed that all lamin constructs were expressed at similar levels (Extended Data Figure 2) and correctly localized to the nucleus following dox treatment (Figure 1F-H).

Micropipette aspiration of these cell lines revealed that dox induced expression of either A-type lamin (LaA or LaC) in *Lmna*^-/-^ MEFs significantly reduced nuclear deformation compared to the vehicle-treated (‘no dox’) baseline controls (Figure 1I-J). In contrast, LaB1 overexpression did not change nuclear deformability in *Lmna*^-/-^ MEFs (Figure 1K), suggesting that LaB1 cannot overcome the mechanical defects associated with loss of A-type lamins, whereas either LaA or LaC is sufficient to rescue these defects to a similar extent. In TKO MEFs, expression of LaA restored nuclear deformability to wild-type levels (Figure 1M), i.e., achieved full rescue, even in the absence of B-type lamins. Expression of LaC also substantially reduced nuclear deformability of TKO cells, albeit to a lesser degree than LaA (Figure 1M). The more limited rescue of LaC compared to LaA may be due to the higher expression levels of LaA in TKO MEFs, whereas in *Lmna*^-/-^ MEFs, the expression levels of LaA and LaC were closely matched and resulted in similar nuclear stiffness (Figure 1I, J). Whereas expression of LaB1 did not alter nuclear stiffness in *Lmna*^-/-^ MEFs, LaB1 expression in TKO MEFs significantly reduced nuclear deformability (Figure 1O), albeit to a lesser extent than LaA expression. Notably, LaB1 expression restored nuclear deformability of TKO cells to levels observed in *Lmna*^-/-^ MEFs (Figure 1C, D), which lack lamin A/C but have normal levels of lamin B1 and B2^19^. These findings indicate that expression of LaB1 is sufficient to rescue the loss of both B-type lamins, further supporting the idea that both LaB1 and LaB2 play similar functions in nuclear mechanical stability. Collectively, these results indicate that both A- and B-type lamins modulate nuclear stiffness, but to different extents, since LaA could completely compensate for loss of all lamins, whereas overexpression of LaB1 could not overcome loss of A-type lamins in the *Lmna*^-/-^ MEFs, and LaB1 expression had only a moderate effect on the deformability of TKO cells.

### A-type and B-type lamins both contribute to the stiffness and viscosity of the nuclear interior

The lamin-specific differences in rescuing the nuclear stiffness of *Lmna*^-/-^ and TKO MEFs could arise from distinct effects of specific lamins on biophysical and biochemical factors that modulate nuclear mechanics, including (1) altering physical chromatin organization and stiffness, e.g., by cross-linking chromatin or affecting chromatin compaction^41,43^, (2) tethering the nuclear interior to the nuclear membrane, which increases nuclear stiffness^44^, (3) biochemical or structural differences in the head-, rod-, or tail-domain between the different lamin types that alter the physical stability of the lamin network^13^, or (4) differential interactions with binding partners that contribute to nuclear stiffness^45^. Since loss of A-type lamins is known to affect chromatin organization^46^, which could alter nuclear stiffness^25,43^, we first investigated the effect of specific lamins on the physical properties of the nuclear interior, using two complementary non-invasive techniques: Brillouin microscopy and diffusivity of nuclear genetically encoded multimeric nanoparticles (nucGEMs).

Brillouin microscopy is a label-free method to probe the local mechanical properties of materials based on light scattering, where interaction of incident light with acoustic phonons within a material causes a frequency shift in the scattered light^47^. Brillouin microscopy has high spatial resolution and has recently emerged as a useful technique to measure the mechanical properties of subcellular structures, such as the nucleus, in live, adherent cells^48–52^. We hypothesized that loss of lamins results in reduced chromatin stiffness, corresponding to a decreased Brillouin frequency shift, due to reduced intranuclear chromatin cross-linking by lamins and/or the detachment of lamina-associated chromatin domains from the nuclear envelope^43,44^.

Consistent with this hypothesis, TKO MEFs had a decreased Brillouin frequency shift in the nucleus compared to wild-type controls (Figure 2A-C), suggesting that loss of lamins results in decreased chromatin stiffness. However, expression of either LaA or LaB1 in TKO MEFs, representing A- and B-type lamins, respectively, restored the Brillouin frequency shift to wild-type levels (Figure 2D-E). These findings indicate that both A- and B-type lamins are sufficient to rescue chromatin mechanics associated with loss of all lamin proteins. Thus, the lamin-specific effects observed in our micropipette aspiration assay cannot be explained by differential effects of specific lamin types on chromatin organization and mechanics.

**Figure 2:**
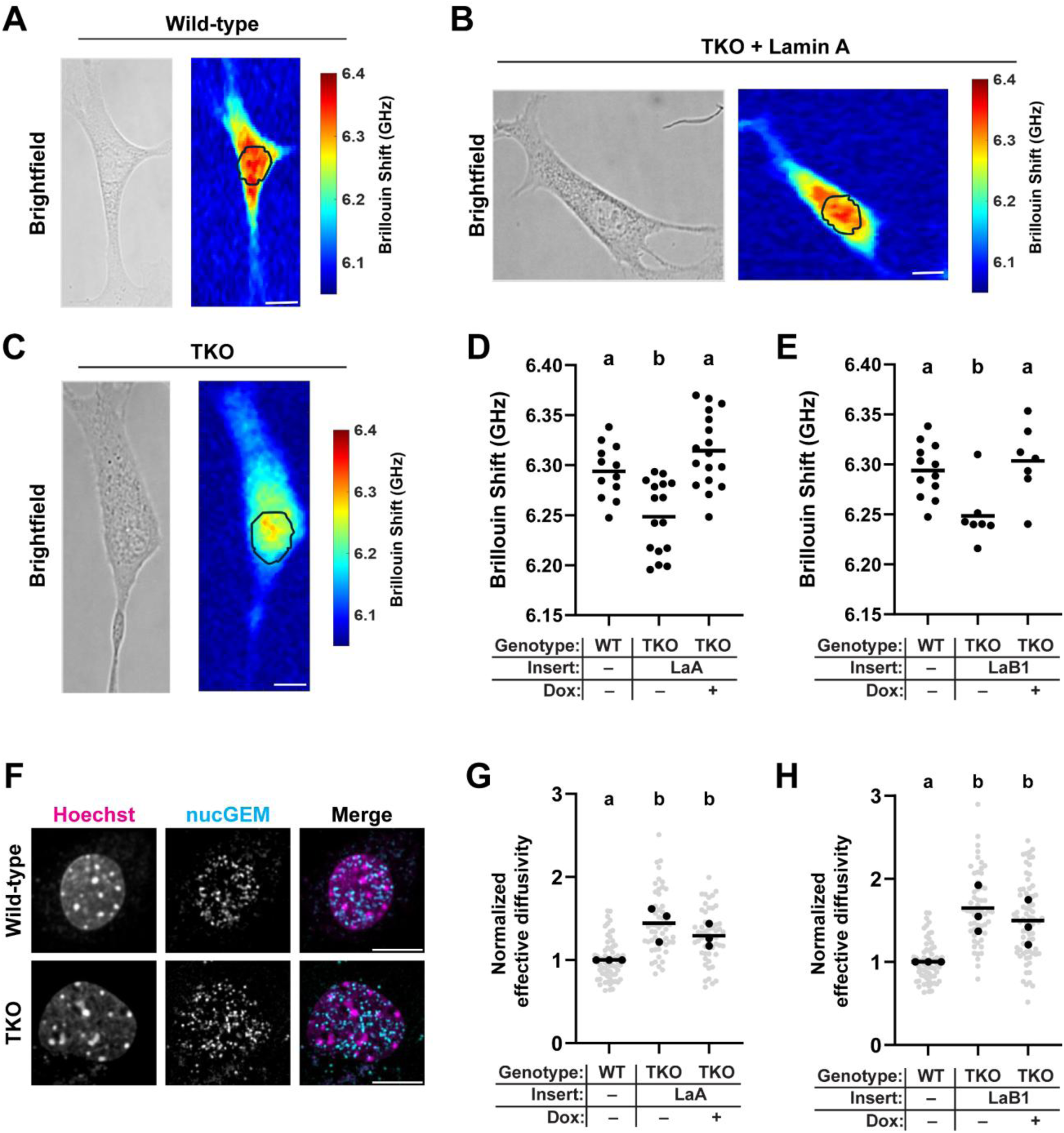
Lamins influence viscoelastic properties of the nuclear interior. (**A-C**) Representative images of WT, TKO, or TKO + LaA rescue cells. Brightfield images show the boundaries of cells, and the Brillouin frequency shift is pseudocolored to depict the spatial distribution of Brillouin shift across the cell. The nucleus is marked by the black outline. Scale bar = 10 µm. (**D-E**) Quantification of Brillouin frequency shift for cells of the indicated genotype and treatment conditions. Points represent measurements from individual cells, bars indicate means. Different letters above each set of points represent *p* < 0.05 based on One-way ANOVA with Tukey’s multiple comparison test. (**F**) Representative images of wild-type or TKO MEFs stably modified to express nucGEMs. Scale bar = 10 µm. (**G-H**) Quantification of normalized nucGEM effective diffusivity, relative to the mean wild-type value for each replicate. Grey points indicate measurements from individual cells, black points indicate replicate means, and bars indicate the overall mean.

As an orthogonal approach to examine the contributions of specific lamin types to the mechanical properties of chromatin, we used nucGEMs, a recently developed tool to probe the mesoscale biophysical properties of the nuclear interior in live cells^53,54^. These self-assembling nanoparticles, each ∼50 nm in diameter, are composed of fluorescently tagged *Quasibacillus thermotolerans* encapsulin proteins (which do not interact with endogenous proteins) fused to a nuclear localization signal, causing them to localize to the nuclear interior (Figure 2F). High-resolution tracking of the nucGEM nanoparticles can be used to compute their effective diffusivity, which is influenced by local crowding, elastic confinement, and viscosity. TKO MEFs exhibited significantly larger effective diffusivity, corresponding to increased mobility of nucGEMs, compared to wild-type controls (Figure 2G-H). However, neither expression of LaA nor expression of LaB1 in TKO MEFs significantly reduced effective diffusivity (Figure 2G-H), suggesting that a combination of different lamin types is necessary to influence the mesoscale fluidity measured by nucGEM mobility. We postulate that lamins jointly modulate the biophysical properties of the nuclear interior, since nucGEMs were more mobile in cells lacking all lamins, but that the experimental assay lacks the sensitivity to evaluate the contributions of specific lamins to nuclear fluidity. Taken together, both the Brillouin and nucGEM measurements suggest that lamins contribute to the mechanical properties of the nuclear interior, but neither approach revealed differences between the A-type and B-type lamins. Thus, these data suggest that the lamin-specific differences in nuclear stiffness observed by the micropipette aspiration do not arise from A-type vs B-type lamin-specific effects on chromatin mechanics, but instead from differences in the assembled lamin networks at the nuclear periphery.

### Anchoring to the nuclear membrane via farnesyl enhances the nuclear stiffness effect of lamin A, but not lamin B1

One major difference between A- and B-type lamins is their anchoring to the inner nuclear membrane via a C-terminal farnesyl group, which has been previously proposed as an explanation for the divergent functions of LaA and LaB1^7,55^. During normal lamin processing, LaA and B-type lamins (but not LaC) are farnesylated at the C-terminal cysteine in the CaaX motif, where ‘C’ represents a cysteine, ‘a’ an aliphatic amino acid, and ‘X’ any amino acid^56^. However, whereas B-type lamins retain this farnesylation, farnesylated prelamin A undergoes a proteolytic cleavage of its C-terminal 15 amino acids, including the farnesylated CaaX motif, resulting in mature LaA that is not farnesylated. Since permanent farnesylation of mutant LaA, as seen in Hutchinson-Gilford Progeria Syndrome (HGPS), results in increased nuclear stiffness^57–59^, we examined how modulating farnesylation of LaA and LaB1 affected their contribution to nuclear stiffness using our exogenous, inducible expression system in the *Lmna*^-/-^ and TKO MEFs. Because the rapid processing of prelamin A into mature LaA precludes performing experiments with permanently farnesylated LaA, we expressed progerin, a permanently farnesylated form of mutant LaA that is responsible for HGPS and that contains a 50 amino acid deletion, which includes the cleavage recognition motif^60^. Progerin expression in *Lmna*^-/-^ MEFs resulted in wrinkled nuclei that were substantially less deformable than those in cells expressing wild-type LaA (Figure 3A-D).

**Figure 3:**
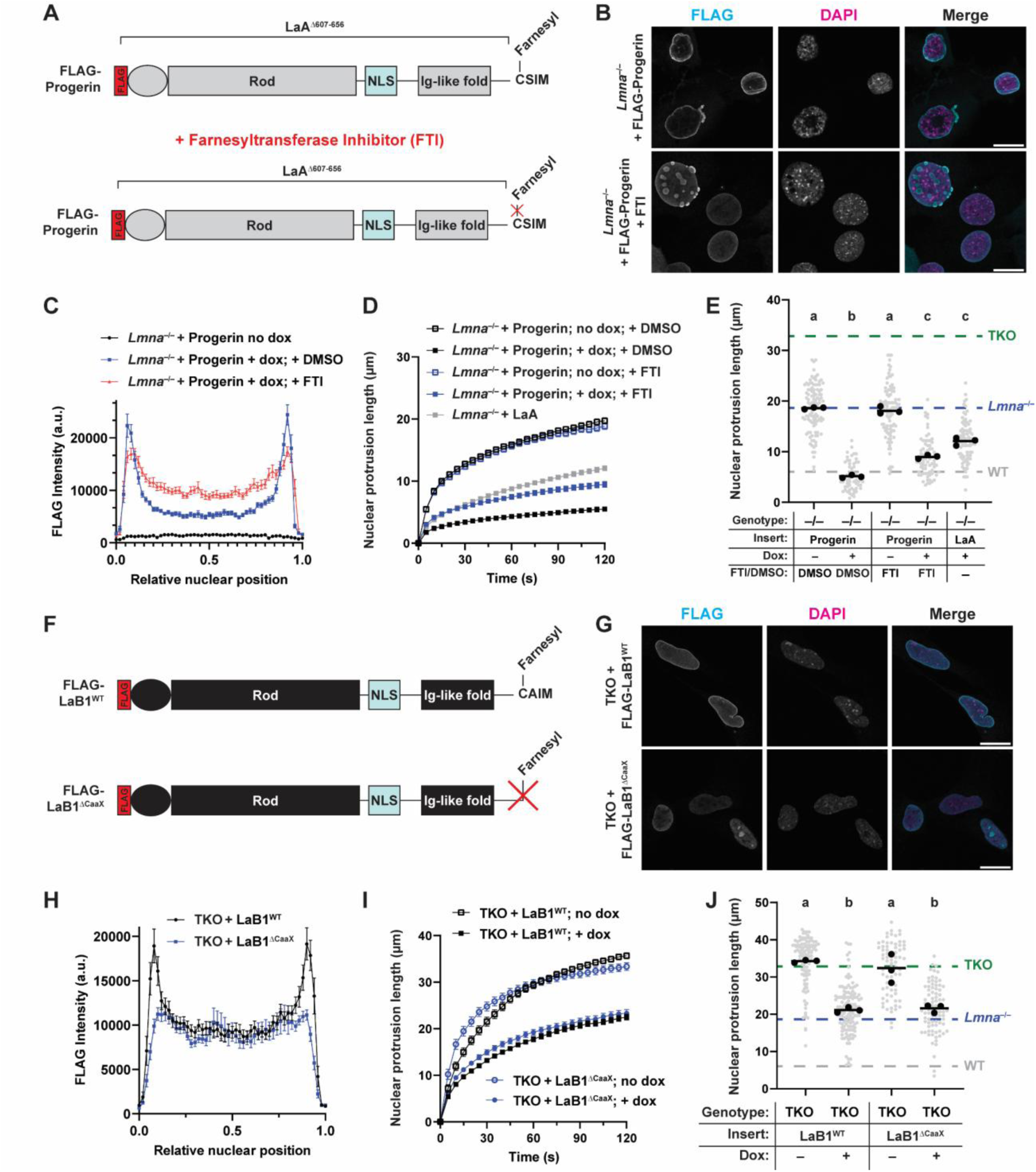
Tethering lamins to the nuclear membrane via farnesylation increases nuclear stiffening by LaA but not LaB1. (**A**) Schematic depiction of FLAG-Progerin expression construct, which lacks amino acids 607-656 of LaA. The addition of a farnesyltransferase inhibitor (FTI) blocks farnesylation of this protein. (**B**) Representative images of *Lmna*^-/-^ MEFs expressing progerin, treated with vehicle (DMSO) or FTI. Treatment with FTI blocks farnesylation of progerin, resulting in a loss of signal at the nuclear periphery. Scale bar = 20 µm. (**C**) FLAG intensity profiles across a line drawn through the midplane of the nucleus. Treatment with FTI results in an increase in the nucleoplasmic signal. (**D-E**) Quantification of nuclear deformability over time of *Lmna*^-/-^ MEFs expressing progerin, treated with either DMSO or FTI, or LaA as control. (**F**) Schematic of wild-type or CaaX deleted (ΔCaaX) LaB1 constructs. Deletion of the CaaX motif (CAIM) prevents farnesylation. (**G**) Representative confocal images through the midplane of the nucleus of TKO MEFs expressing wild-type or CaaX-deleted LaB1. Scale bar = 20 µm. (**H**) anti-FLAG immunofluorescence intensity profiles across a line drawn through the midplane of the nucleus. Peaks at the edges represent the nuclear rim staining present in LaB1^WT^, and the signal in the center represents the nucleoplasmic LaB1. Points represent the average of 25-30 cells, error bars represent the overall s.e.m. (**I-J**) Quantification of nuclear deformability of TKO MEFs expressing LaB1^WT^ or LaB1^ΔCaaX^. Grey points indicate measurements from individual cells, black points indicate replicate means, and bars indicate overall means. Sets of points with the same letter above them are not significantly different, whereas different letters indicate *p* < 0.05, based on One-way ANOVA with Tukey’s multiple comparison test.

Treatment with a farnesyltransferase inhibitor (FTI), lonafarnib, which prevents farnesylation of progerin^61^, led to re-localization of progerin from the nuclear periphery to the nuclear interior (Figure 3B-C), consistent with a lack of anchoring to the nuclear membrane. FTI treatment reduced the nuclear deformability of *Lmna*^-/-^ MEFs expressing progerin to levels comparable to those of *Lmna*^-/-^ MEFs expressing wild-type LaA (Figure 3D-E), indicating that the additional increase in stiffness associated with progerin expression was due to the farnesylation and not the 50-amino acid deletion of progerin. These results suggest that farnesylation and anchoring of LaA to the nuclear membrane increases its effect on nuclear stiffness.

To study the effect of farnesylation on LaB1, we generated TKO MEFs with inducible expression of a LaB1 construct lacking the CaaX motif, LaB1^ΔCaaX^, which prevents farnesylation. Deletion of the CaaX motif resulted in relocalization of LaB1^ΔCaaX^ from the nuclear periphery to the nuclear interior in the TKO MEFs (Figure 3F-H), similar to the effect of FTI treatment on progerin localization. Surprisingly, though, both LaB1^ΔCaaX^ and LaB1 rescued nuclear stiffness in TKO MEFs to similar extent (Figure 3I-J). A CaaX mutated LaB1 construct (LaB1^CAIMS^), which also exhibited defective localization to the nuclear periphery, similarly rescued nuclear deformability of TKO MEFs to the same degree as wild-type LaB1 (Extended Data Figure 3). These results suggest that, unlike the effect seen in LaA, the contribution to nuclear stiffness conferred by LaB1 is not dependent on its farnesylation and its anchoring to the inner nuclear membrane. Furthermore, these results suggest that farnesylation cannot account for the difference in the A-type versus B-type lamin-specific contributions to nuclear stiffness.

### The larger contribution of LaA to nuclear stiffness is conferred by its rod domain

To determine the basis for these lamin-specific contributions to nuclear stiffness, we compared the primary amino acid sequence of human LaA and LaB1 (Extended Data Figure 1). We found that the proteins are most divergent in their tail domains, which contain the Ig-like fold. Therefore, we aimed to determine whether the more substantial effect of LaA in providing mechanical stability to the nucleus was due to its tail domain, which is responsible for many protein-protein interactions^62^, or due to its head and rod domains. We generated different truncated lamin constructs that consisted either of each protein’s head and rod, or the tail (Figure 4A). These truncation constructs were designed so that each retained the original nuclear localization sequence (NLS) to ensure proper targeting to the nucleus (Figure 4B). However, unlike the full-length lamin constructs, the truncated lamins had a strong nucleoplasmic presence and were at most only slightly enriched at the nuclear periphery, unlike the full-length lamin constructs (compare Figure 4B with Figure 1F-H), suggesting that the full-length sequence is necessary for proper incorporation and/or assembly of lamins into the lamina. We did not observe any differences in the subcellular localization of the LaA^Head+Rod^ and LaB1^Head+Rod^ truncations, nor in their solubility, as assessed by differential protein extraction using different stringency buffers (Extended Data Figure 2).

**Figure 4:**
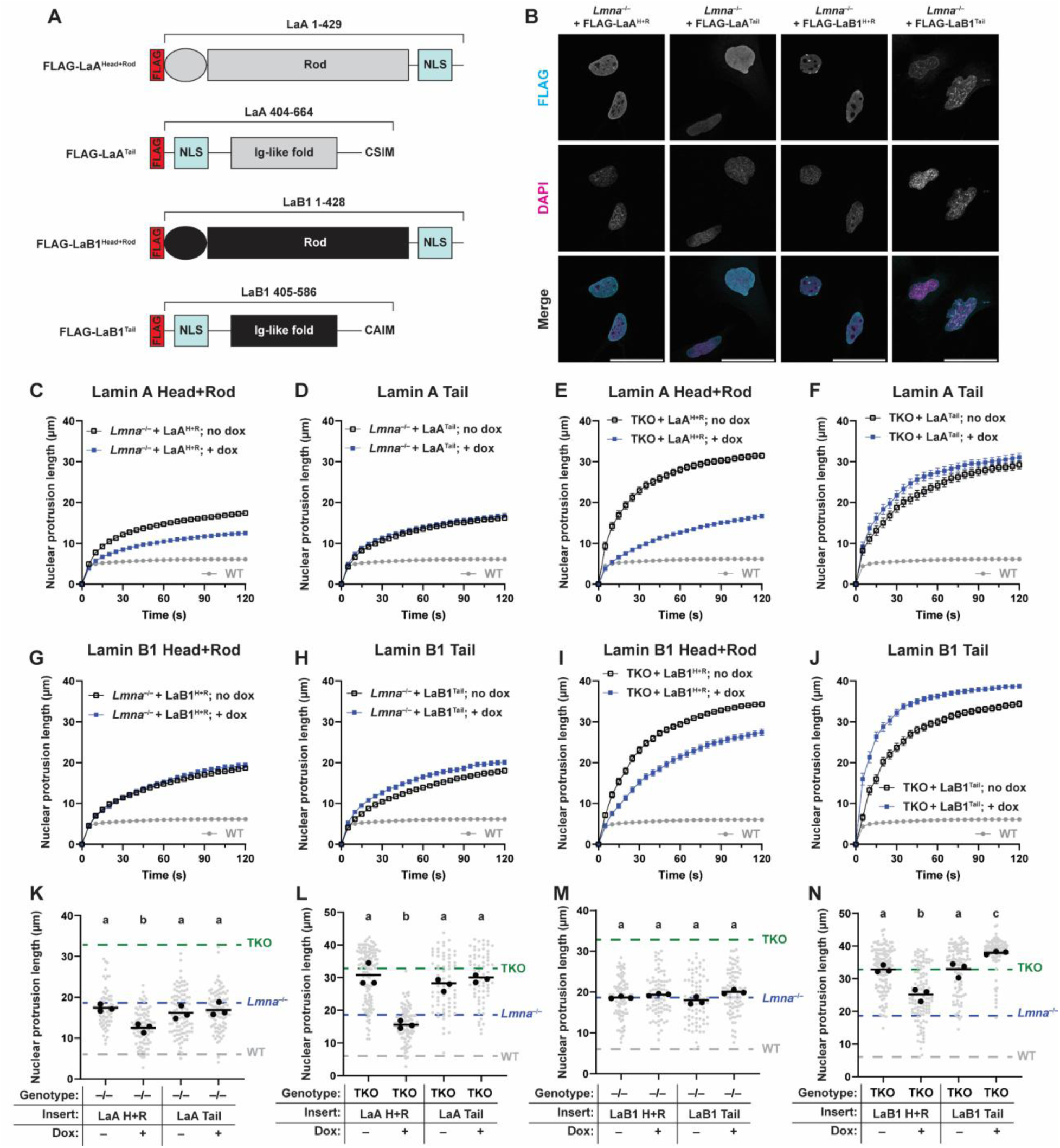
Lamin Head+Rod domains differentially regulate nuclear deformability. (**A**) Schematic representation of the different lamin truncations, with the amino acids from LaA or LaB1 indicated. (**B**) Representative immunofluorescence staining of *Lmna*^-/-^ MEFs expressing truncated lamin constructs. Scale bar = 20 µm. (**C-F**) Time-dependent nuclear protrusion of *Lmna*^-/-^ MEFs (**C-D**) or TKO MEFs (**E-F**) expressing LaA^Head+Rod^ or LaA^Tail^. (**G-J**) Time-dependent nuclear protrusion of *Lmna^-/-^* (**G-H**) or TKO MEFs (**I-J**) expressing LaB1^Head+Rod^ or LaB1^Tail^. (**K-N**) Quantification of nuclear protrusion length of cells expressing LaA truncations (**K-L**) or LaB1 truncations (**M-N**) after 100 s of aspiration. Grey points indicate measurements from individual cells, black points indicate replicate means, and bars indicate overall means. Sets of points with the same letter above them are not significantly different, whereas different letters indicate *p* < 0.05, One-way ANOVA with Tukey’s multiple comparison test.

Remarkably, the LaA truncation consisting of the head and rod domain (LaA^Head+Rod^) nonetheless significantly reduced nuclear deformability when expressed in *Lmna*^-/-^ and TKO MEFs, mirroring the rescue achieved by full-length LaA and LaC (Figure 4C, E, K, L). The truncation containing the head and rod of LaB1 (LaB1^Head+Rod^) did not significantly alter nuclear deformability when expressed in *Lmna*^-/-^ MEFs (Figure 4G, M) but led to a significant reduction in deformability when expressed in TKO MEFs (Figure 4I, N), mirroring the rescue observed with full-length LaB1 (compare with Figure 1O). In contrast to the truncation containing the head and rod domains, neither Ig-Fold containing truncation (LaA^Tail^ and LaB1^Tail^) reduced nuclear deformability when expressed in *Lmna*^-/-^ or TKO MEFs. In fact, the LaB1^Tail^ construct caused a slight *increase* in nuclear deformability when expressed in TKO MEFs (Figure 4J, N) along with abnormally shaped nuclei (Figure 4B), possibly by increasing nuclear surface area or causing the mislocalization of other inner nuclear membrane proteins such as LAP1, as previously reported for a LaB1 truncation lacking the rod domain^63^.

Brillouin and nucGEM experiments performed using TKO cells expressing the LaA^Head+Rod^ and LaB1^Head+Rod^ truncations did not reveal any differences between the A-type and B-type constructs (Extended Data Figure 4), indicating that despite the large nucleoplasmic presence of the lamin truncation, the LaA specific effect on nuclear stiffness is likely due to its effect at the nuclear periphery. Taken together, our results indicate that the divergent ability of A-type and B-type lamins to stiffen nuclei is mediated by residues found in the head and rod domains of the respective proteins, rather than their Ig-fold, and is likely linked to differences in their filament formation or filament properties.

To further confirm the importance of the head and tail domain of LaA in conferring nuclear stiffness, we generated chimeric lamin constructs that combined either the head and rod domain of LaA with the tail of LaB1 (LaA^Head+Rod^-LaB1^Tail^) or the head and rod domain of LaB1 with the tail of LaA (LaB1^Head+Rod^-LaA^Tail^), and expressed these chimeras in *Lmna*^-/-^ and TKO MEFs. Expression of the LaB1^Head+Rod^-LaA^Tail^ construct closely resembled the limited rescue of nuclear stiffness achieved by LaB1, rather than the more complete rescue by LaA (Extended Data Figure 5). In contrast, expression of LaA^Head+Rod^-LaB1^Tail^ in *Lmna*^-/-^ MEFs resulted in nuclei that were wrinkled and very stiff (Figure 5A-G), resembling the effect of progerin expression (see Figure 2), which, like the tail of LaB1, is also permanently farnesylated. Similar to the stiffening effect associated with progerin expression, the stiffness conferred by the LaA^Head+Rod^-LaB1^Tail^ construct was also partially dependent on farnesylation. FTI treatment led to a more intranuclear localization of this protein (Figure 5B-C) and partially reversed the stiffness conferred by this construct in *Lmna*^-/-^ and TKO MEFs (Figure 5D-G). To test whether any anchoring of the LaA head and rod domain to the inner nuclear membrane resulted in progerin-like increases in nuclear stiffness, we generated an additional chimera that combined the head and rod of LaA with the tail of the distantly related amoeba lamin homolog, NE81, which is also permanently farnesylated^42,64^. Expression of the LaA^Head+Rod^-NE81^Tail^ chimera in *Lmna*^-/-^ and TKO MEFs resulted in wrinkled, stiff nuclei; the effect was partially reversed with FTI treatment (Figure 5H-N), similar to the results seen for progerin and the LaA^Head+Rod^-LaB1^Tail^ chimera. Thus, the combination of the head and rod of LaA with any permanently farnesylated lamin tail results in progerin-like effects on nuclear stiffness, implicating the LaA head and rod domains as a key driver for the behavior of these chimeric lamins.

**Figure 5:**
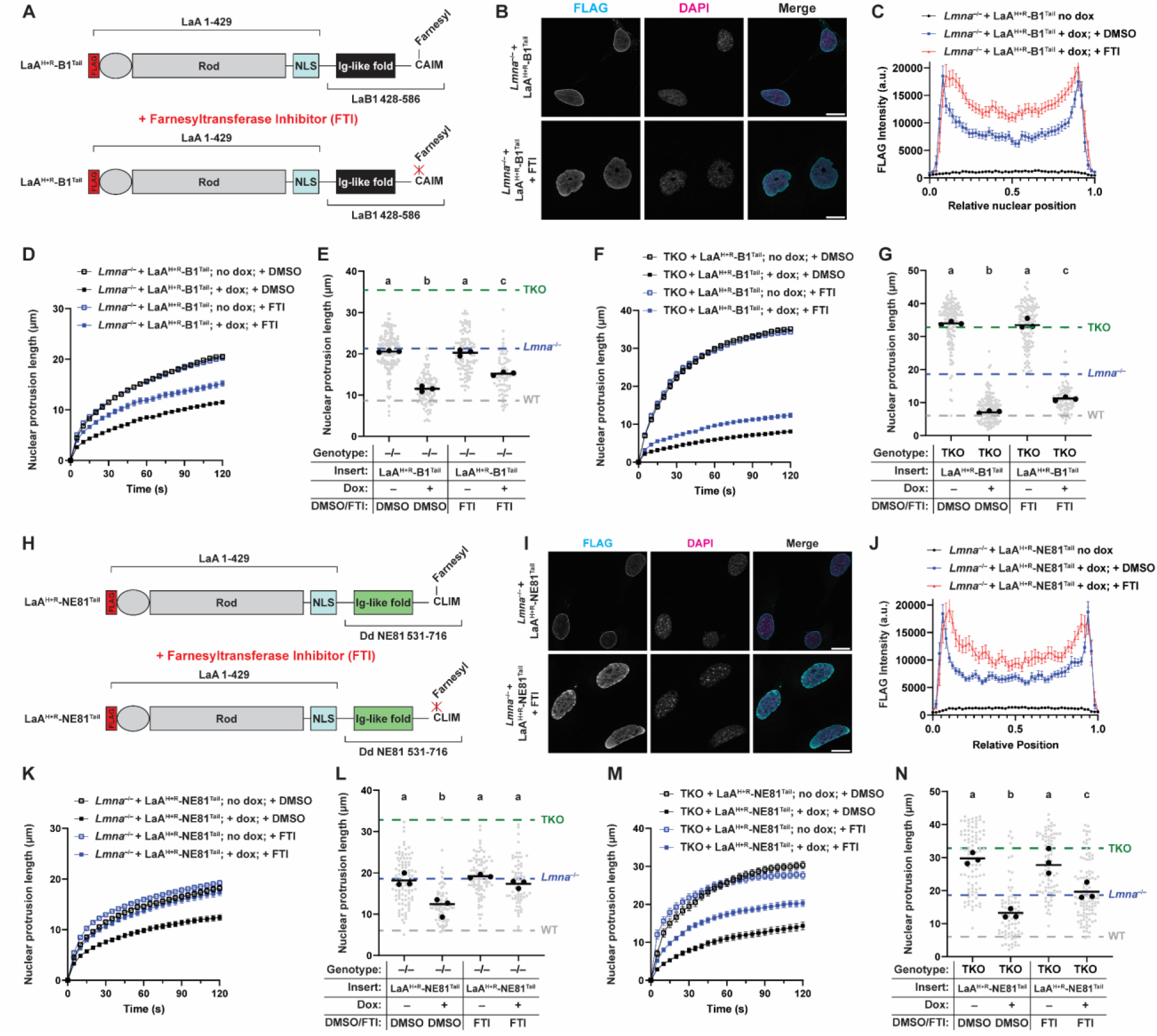
Tethering LaA Head+Rod to the INM stiffens nuclei in a farnesylation-dependent manner. (**A**) Schematic of LaA^Head+Rod^-LaB1^Tail^ expression construct, which combines the head and rod of LaA with the tail of LaB1. (**B**) Representative images of *Lmna*^-/-^ MEFs expressing this construct, treated with vehicle (DMSO) or FTI. Scale bar = 20 µm. (**C**) FLAG intensity profiles across a line drawn through the midplane of the nucleus. Treatment with FTI results in an increase in LaA^Head+Rod^-LaB1^Tail^ nucleoplasmic signal. (**D-G**) Quantification of nuclear deformability of *Lmna^-/-^*MEFs (**D-E**) or TKO MEFs (**F-G**) expressing LaA^Head+Rod^-LaB1^Tail^, treated with either DMSO or FTI. (**H**) Schematic of LaA^Head+Rod^-NE81^Tail^ expression construct, which combines the head and rod of LaA with the tail of the *D. discoideum* lamin homolog NE81. (**I**) Representative images of *Lmna*^-/-^ MEFs expressing this construct, treated with vehicle (DMSO) or FTI. Scale bar = 20 µm. (**J**) FLAG intensity profiles across a line drawn through the midplane of the nucleus. Treatment with FTI results in an increase in LaA^Head+Rod^-NE81^Tail^ nucleoplasmic signal. (**K-N**) Quantification of nuclear deformability of *Lmna^-/-^* MEFs (**K-L**) or TKO MEFs (**M-N**) expressing LaA^Head+Rod^-NE81^Tail^, treated with either DMSO or FTI. Grey points indicate measurements from individual cells, black points indicate replicate means, and bars indicate overall means. Sets of points with the same letter above them are not significantly different, whereas different letters indicate *p* < 0.05, One-way ANOVA with Tukey’s multiple comparison test.

Given the high degree of similarity between the lamin rod domains at the amino acid level, we considered whether the A-type versus B-type lamin-specific differences in rescue ability observed in our lamin truncations and chimeras might be due to differences in the small N-terminal head domains, which differ substantially between A-type and B-type lamins (Extended Data Figure 1). The heads of lamins and other intermediate filaments are critical for proper assembly of lamins into higher-order structures, both of which are necessary to confer mechanical strength to nuclei^4,65^. To test whether the identity of the head domain could explain the differences between A- and B-type lamins, we generated lamin constructs that swapped the head domains of full-length LaA and LaB1, as well as head-swapped truncations that lacked the lamin tail domains (Figure 6A). All the head-swapped lamins localized to the nucleus as expected (Figure 6B) and mirrored the distribution of their “wild-type” counterparts, except the LaB1^Head^-LaA^Rod+Tail^ construct, which frequently formed punctate structures at the nuclear periphery (Figure 6B).

**Figure 6:**
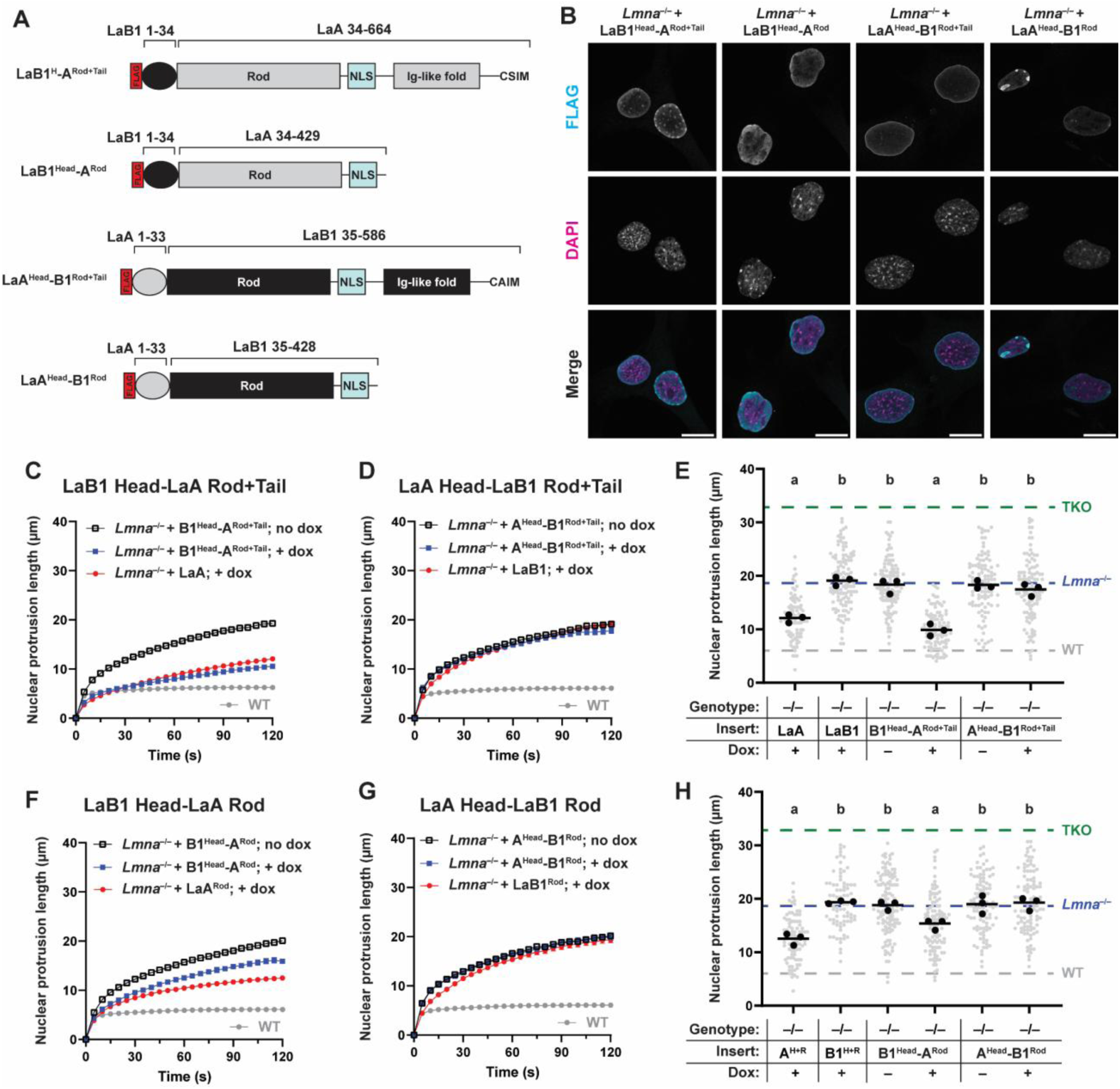
The identity of the lamin rod domain determines the stiffening ability of lamins. (**A**) Schematic of ‘head-swapped’ lamin constructs, with the residues from LaA or LaB1 indicated. (**B**) Representative immunofluorescence staining of *Lmna*^-/-^ MEFs expressing head-swapped lamin constructs. Scale bar = 20 µm. (**C-E**) Time-dependent nuclear protrusion of *Lmna*^-/-^ MEFs expressing LaB1^Head^–LaA^Rod+Tail^ (**C**) or LaA^Head^–LaB1^Rod+Tail^ (**D**), with quantification of nuclear protrusion length following 100 s of aspiration (**E**). Data from *Lmna*^-/-^ MEFs expressing LaA or LaB1 are shown for reference. (**F-H**) Time-dependent nuclear protrusion of *Lmna*^-/-^ MEFs expressing LaB1^Head^ –LaA^Rod^ (**F**) or LaA^Head^ –LaB1^Rod^ (**G**), with quantification of nuclear protrusion length following 100 s of aspiration (**H**). Data from *Lmna*^-/-^ MEFs expressing LaA^Head+Rod^ or LaB1^Head+Rod^ are shown for reference. Grey points indicate measurements from individual cells, black points indicate replicate means, and bars indicate overall means. Sets of points with the same letter above them are not significantly different, whereas different letters indicate *p* < 0.05, based on One-way ANOVA with Tukey’s multiple comparison test.

Despite this unusual localization, the addition of the LaB1 head to the LaA rod and tail did not reduce the rescue ability of the LaB1^Head^-LaA^Rod+Tail^ construct, and the nuclear deformability of *Lmna* ^-/-^ or TKO MEFs expressing LaB1^Head^-LaA^Rod+Tail^ was not significantly different from those expressing full-length LaA (Figure 6C; Extended Data Figure 6). A truncated lamin chimera, consisting of the LaB1 head and the LaA rod but not the LaA tail (LaB1^Head^-LaA^Rod^), was sufficient to rescue nuclear stiffness when expressed in *Lmna*^-/-^ or TKO MEFs; the LaB1^Head^-LaA^Rod^ chimera achieved similar rescue of nuclear stiffness as the LaA^Head+Rod^ truncation (Figure 6D-E, Extended Data Figure 6). In contrast, reciprocal constructs that combined the LaA head with the rod or the rod and tail domains of LaB1 did not improve the rescue achieved by LaB1 (Figure 6F-H; Extended Data Figure 6), indicating that the LaA head domain is not responsible for the increased nuclear stiffening effect of LaA. Taken together, our data demonstrate that the identity of the lamin rod domain determines the divergent contributions to nuclear mechanics of the A-type and B-type lamins, with the A-type lamin rod domain conferring a stronger effect on nuclear stiffness than the LaB1 rod domain.

### Identification of A-type vs B-type lamin-specific interaction partners

Previous in vitro assembly studies found that truncations containing the head and rod domains of lamins were sufficient to form filaments^4,5^. Thus, the strong effect of the LaA rod on nuclear stiffness could arise from the mechanical properties of the resulting filaments or from specific interactions between LaA and other proteins that confer increased nuclear stability. Since the mechanical properties of lamin filaments are not accessible for direct experimental measurements, except for in the large *Xenopus* oocytes, which have a unique lamin isoform expression and network architecture^66^, we focused on identifying specific interaction partners of LaA vs LaB1.

Although extensive prior work has identified numerous interaction partners of specific lamin proteins via proximity-based approaches such as Bio-ID^67,68^ or affinity-based techniques^62,69^, no direct comparison of the interaction partners of A-type and B-type lamins has been reported to date. Therefore, we decided to take advantage of the unique opportunity to directly compare interactomes of full-length lamins and lamin truncations by using the N-terminal FLAG tag of our expression construct as bait for co-immunoprecipitation (co-IP) experiments. Co-IPs were performed as outlined in Figure 7A using full-length LaA, LaB1, and the respective truncations consisting of only the head and rod domains, expressed in *Lmna*^-/-^ MEFs. Following co-IP, using each construct as the bait protein, we identified affinity-based interactors by mass spectrometry, comparing each IP with a negative control IP performed on the parental *Lmna*^-/-^ MEFs (Figure 7B; Extended Data Figure 7).

**Figure 7:**
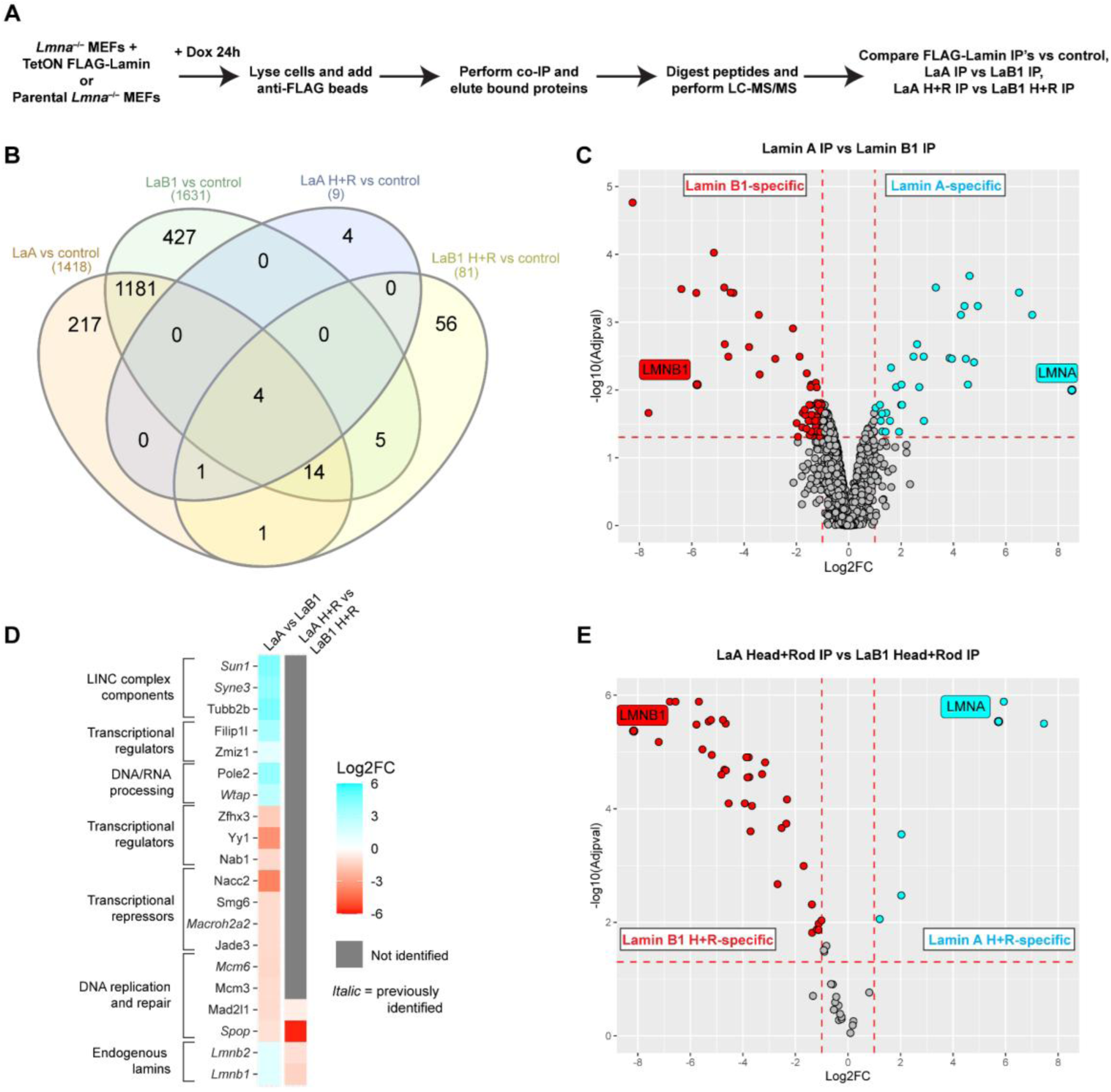
Identification of LaA versus LaB1-specific interaction partners by co-IP mass spectrometry. (**A**) Outline of co-IP mass spec experiments. (**B**) Venn diagram comparison of significantly enriched interactors of each bait protein. Proteins included in each list were identified by comparing the proteins identified using each bait versus a negative control IP (for individual volcano plots, see Extended Data 7). (**C**) Volcano plot comparing the interaction partners of LaA versus LaB1. Positive values of log2-fold-change (log2FC) indicate enrichment in the LaA co-IP, whereas negative values indicate enrichment in the LaB1 co-IP. Proteins that had a log2FC greater than 1 or less than -1 and had an adjusted p-value < 0.05 are color-coded as preferentially interacting with LaA (cyan) versus LaB1 (red), respectively. The bait proteins (human LaA/LMNA and LaB1/LMNB1) are indicated. (**D**) Selected lamin-specific interaction partners, based on comparison performed in (**C**) and (**D**). Interactors were manually grouped by shared function/processes, and proteins in italics have been previously identified to interact with lamins (either mouse or human). Grey coloring indicates that the protein was not identified in the co-IP performed using the rod truncations, and thus there is no enrichment in one IP versus the other. (**E**) Volcano plot depicting the comparison of interaction partners of the lamin head+rod truncations, as in (**C**).

Using the full-length LaA and LaB1 constructs, we identified numerous interactions shared between full-length LaA and LaB1, and also many that appeared only in one co-IP and not the other (217 for LaA and 427 for LaB1). Of all candidate interactors identified, 159 (14%) and 99 (7%) had been previously determined to interact with LaA or LaB1, respectively. Direct comparison between the co-IPs performed on LaA versus LaB1 yielded a more stringent list of A-type versus B-type specific interaction partners (Figure 7C), with selected proteins shown in Figure 7D. This list included previously identified lamin binding partners, such as Sun1, which interacts specifically with LaA^70^, and novel candidate interactors, such as the Filamin A-interacting protein 1-like, Filip1l, a tumor-suppressor-like protein that inhibits canonical WNT signaling^71^. We validated the preferential interaction of Filip1l with LaA by co-IP (Extended Data Figure 8A-B). Subcellular localization of Filip1l, however, did not depend on lamins (Extended Data Figure 8C), so the functional significance of this interaction remains to be determined.

In contrast to the co-IP mass spec analysis performed on full-length lamin constructs, co-IPs using the LaA^Head+Rod^ and LaB1^Head+Rod^ constructs yielded substantially fewer interaction partners (a total of 9 for LaA^Head+Rod^ and 81 for LaB1^Head+Rod^), despite similar enrichment of the FLAG-tagged lamin constructs relative to the control (Supplemental Data Figure 1). These results are consistent with previous reports that the lamin tail domain, and in particular the Ig-like fold, is a hub for interaction with other proteins^62,72^. Direct comparison of the interactors identified using LaA^Head+Rod^ vs LaB1^Head+Rod^ as bait identified very few specific interaction partners of the rod truncations (Figure 7E), and none of the LaA-specific proteins identified with full-length LaA as bait were found when using LaA^Head+Rod^ as the bait, emphasizing that LaA-specific interaction partners are likely mediated through the lamin tail domain rather than the head and rod.

Based on these data, we were unable to identify unique interaction partners of the LaA^Head+Rod^ vs LaB1^Head+Rod^ that could explain their differing contributions to nuclear stiffness, supporting the idea that the differences between the A-type and B-type lamins on nuclear mechanical stability are an emergent property of lamina. Nonetheless, our mass spectrometry analyses provide new insights into the crucial role of the lamin tail domain in mediating many lamin interactions, including several lamin type-specific interactions.

### Lamin rod truncations are equally able to resist cell-intrinsic pulling forces on nuclear pores

Having established that the LaA rod domain is crucial for conferring the unique stiffening effect of LaA on the deformability of the cell nucleus when cells are subjected to large external forces (i.e., during micropipette aspiration or external stretch), we aimed to determine whether we can also detect differences between lamins on the ability of the nuclear envelope to resist cell-intrinsic forces. Here, we focused on the ability of lamins to ensure proper nuclear positioning of nuclear pore complexes (NPCs), whose uniform distribution across the nuclear surface depends on lamins^73^. Previous research had shown that in TKO MEFs, dynein-mediated microtubule pulling forces acting on NPCs during G2 and early M phase of the cell cycle lead to NPCs clustering on one side of the nucleus, adjacent to the centrosome^73^. If at least one lamin is present in cells, nuclei can resist these forces, and NPCs remain evenly distributed around the nuclear periphery^36,73^. In our experiments, expression of either LaA, LaB1, or LaC1 in TKO MEFs was sufficient to rescue NPC distribution, with no detectable differences between the different full-length lamins (Figure 8A-D). Extending this work to the lamin truncation constructs, we found that despite their lack of enrichment at the nuclear periphery, expression of either the LaA^Head+Rod^ or the LaB1^Head+Rod^ truncation completely restored the distribution of NPCs in TKO MEFs to wild-type levels (Figure 8C-D). In contrast, neither the LaA^Tail^ nor the LaB1^Tail^ truncations were able to rescue NPC distribution in TKO MEFs, likely due to their inability to form lamin filaments. The LaB1^Tail^ constructs actually *increased* NPC mislocalization (Figure 8C-D), consistent with the disruptive effect we observed in the micropipette aspiration assay (Figure 4). Taken together, our findings revealed that the head and rod of either lamin type are both necessary and sufficient for proper positioning of NPCs and to resist cell-intrinsic forces on the nucleus. Based on these findings, we postulate that although all lamins contribute some mechanical stability to the nuclear envelope, which is sufficient to protect the nuclear envelope from small cell-intrinsic forces, A-type lamins confer cells with ability to better resist large forces on the nucleus, as is critical in muscle and other mechanically stressed tissues, and that this unique ability of A-type lamins is primarily conferred through their rod domain.

**Figure 8:**
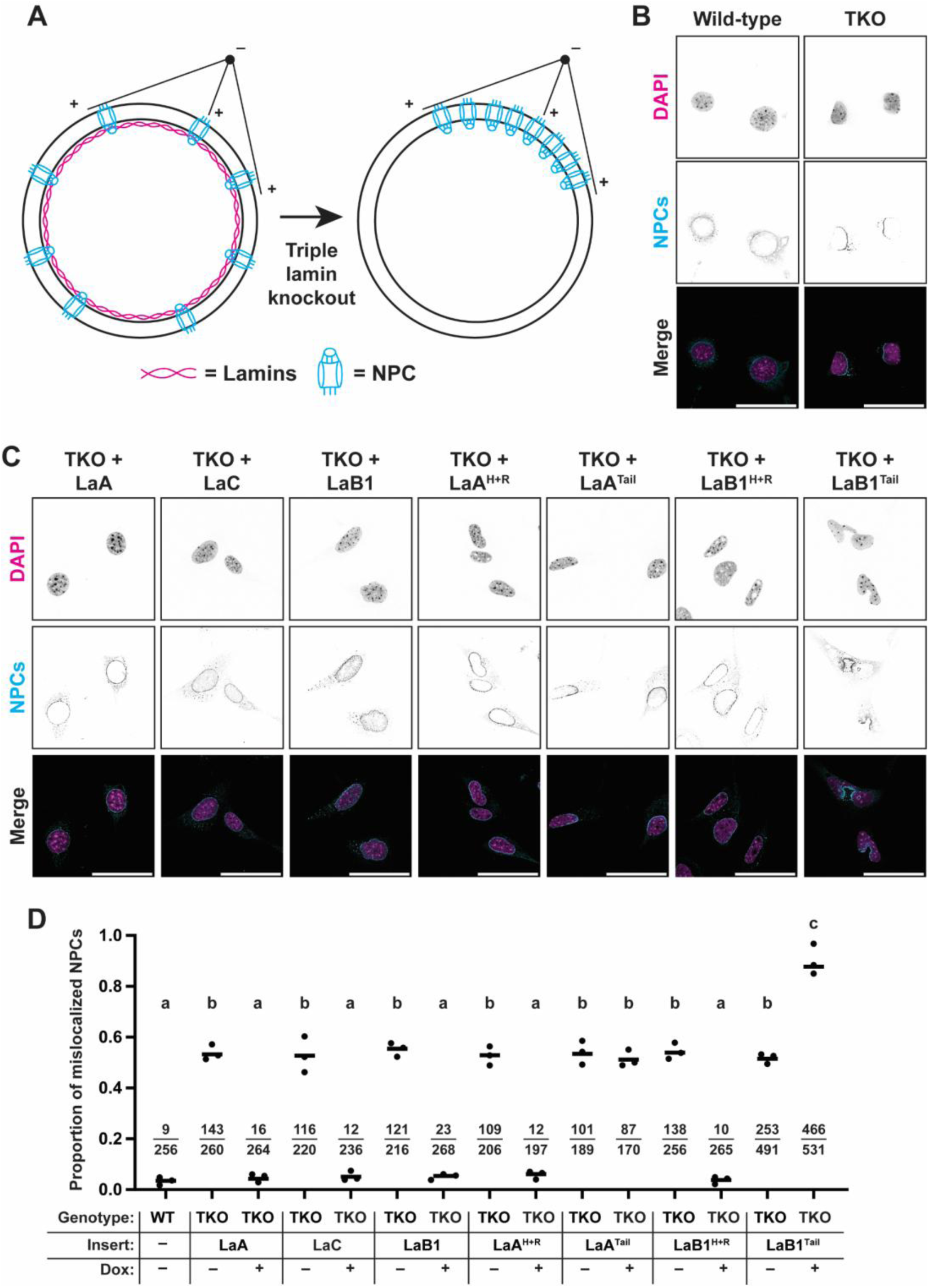
Expression of any individual lamin or Head+Rod truncation complexly rescues NPC positioning defect in TKO MEFs. (**A**) Schematic depiction of microtubule-mediated forces that lead to clustering of NPCs in TKO MEFs (adapted from ref^73^). (**B**) Representative images of wild-type and TKO MEFs stained with the pan-NPC antibody mab414. Scale bar = 50 µm. (**C**) Images of TKO MEFs expressing the various full-length lamins or truncated lamin constructs, stained as in (**B**). Scale bar = 50 µm. (**D**) Quantification of the proportion of nuclei in each condition with mislocalized NPCs, assessed by an observer blinded to the experimental conditions. The fraction of nuclei with ‘mislocalized NPCs’ / ‘total nuclei’ is shown for each condition. Points represent proportions from each of three independent experiments, bars indicate overall proportion pooling all cells. Sets of points with the same letter above them are not significantly different, whereas different letters indicate *p* < 0.05, Fisher’s exact test.

## Discussion

In our study, we aimed to determine the contributions of A-type and B-type lamins to nuclear stiffness and identify the domains responsible for differences between lamin proteins. Using high-throughput micropipette aspiration, we identified significant differences between the ability of A-type and B-type lamins to stiffen nuclei when expressed in lamin-deficient cells and mapped these differences specifically to the lamin rod domain. Surprisingly, differences between the lamin tail domains, which are the most divergent regions of the A-type and B-type lamins, were not sufficient to explain the different contributions to nuclear stiffness between the lamin types. Although LaA and LaB1 differ in their C-terminal anchoring to the INM, the effect of farnesylation depends on whether or not a particular lamin possesses the LaA head and rod or the LaB1 head and rod. We speculate that since the LaA rod has a greater ability to stiffen nuclei compared to the LaB1 rod, tethering of LaA head and rod-containing constructs to the INM via farnesylation results in stiffer nuclei than constructs containing the LaB1 head and rod. This interpretation explains why progerin and LaB1 stiffen nuclei to very different extents, despite both being farnesylated lamins.

Multiple lines of evidence underscore the crucial role of the rod domain in the proper functioning of lamins. Many *LMNA* mutations occur in nucleotides corresponding to the rod domain and are known to destabilize the protein or disrupt filament assembly. Mutations located in the head and rod domain are more likely to result in impaired nuclear stability^24,74–77^. These mutations (e.g., ΔK32, L85R, R60G, N195K) likely prevent the formation of a coiled-coil and assembly into higher order structures^74,76^, but it is unlikely that these residues themselves are sufficient to explain the difference between the A-type and B-type lamins as they are conserved between LaA and LaB1^16,78^. Consistent with the concept that A-type lamins play a crucial role in protecting muscle nuclei from mechanical stress^79^ and our finding that the LaA head and rod are particularly important for this function, mutations in the head and rod domain result in worse disease outcomes for *LMNA*-related dilated cardiomyopathy than those located in the tail domain^80^.

Deletion of the lamin tail domain was previously shown to alter the assembly^4^ and solubility^81^ of lamins, but did not prohibit assembly into higher-order structures. In fact, a rod domain-containing truncation of chicken lamin B2 was capable of assembling into ∼1 μm long filaments in vitro^4^. Subsequent work using truncated lamins found that the N-terminus of one rod domain-containing dimer can interact with the C-terminus of another to form a head-tail tetramer^5^. Given the similarities between these truncations and the ones presented here, we suggest that the lamin constructs used in our studies that consist of the head and rod domains can also assemble into higher-order structures. Lamin assembly has been previously shown to be a prerequisite for nuclear stiffening^82^, and the existence of a stable network is likely required to explain the ability of the lamin truncations to rescue the NPC distribution in lamin-deficient cells.

Other researchers have also examined the contribution of different lamin types to nuclear mechanics. Wintner and colleagues^25^ studied lamin-deficient fibroblasts (*Lmna*^-/-^ and TKO) using a micropipette approach, yet they found only a modest difference in creep compliance between *Lmna*^-/-^ and TKO MEFs, and they reported that expression of either LaA or LaB1 in TKO MEFs resulted in a complete rescue to wild-type levels of deformability, which contrasts with the results presented here. Importantly, the timescales of the measurements differed between our study and theirs–we collected data on nuclear protrusion for 2-3 minutes after aspiration, whereas their study examined nuclear deformation at much shorter timescales (<15 s)^25^. Thus, the differences in our findings may be due to the different viscoelastic regimes being examined. Vahabikashi and colleagues^43^ measured nuclear mechanics by atomic force microscopy (AFM) and optical tweezers (OT) and reported no difference in nuclear stiffness between *Lmna*^-/-^ and *Lmnb1*^-/-^ cells, and that exogenous expression of LaA or LaB1 in *Lmna*^-/-^ cells was equally able to restore wild-type stiffness. However, the authors also note that the deformations caused by AFM or OT are on a relatively small scale, which is a regime that is primarily driven by chromatin states^23^, and indeed, they observed differences in heterochromatin organization upon loss of lamins^43^.

The findings of Vahabikashi et al.^43^ are most consistent with our data collected using Brillouin microscopy, where we observed a decrease in chromatin modulus upon loss of all lamins that was completely rescued by expression of LaA or LaB1. In contrast, the micropipette aspiration system used in our study imposes a large strain on the nucleus, which is primarily resisted by the lamina network^23^. Thus, the specific differences observed with micropipette aspiration are likely due to the emergent lamin network resisting more extensive deformation over a longer timescale. The large deformations applied in our assay over longer times are physiologically relevant, as cells are known to extensively deform their nucleus over minutes to hours during confined migration through tight constrictions or during development^83,84^. Indeed, the balance between A-type and B-type lamins has been repeatedly shown to be an important predictor for cancer cell metastasis and aggressiveness of cancer^85,86^.

Our study contains some limitations. Although we determined that the lamin rod domain determines the stiffening ability of a particular lamin type, we are unable to pinpoint the exact mechanism by which the rods differently contribute to nuclear mechanics. We did not observe obvious differences between the rod truncations’ abilities to modulate chromatin stiffness and were also unable to identify specific interaction partners within the rod domains, suggesting that differential stiffness is an emergent property of the assembled A-type and B-type networks. Future studies could apply a phylogenetic approach to determine highly conserved versus divergent subdomains of the lamin rod across vertebrate A-type and B-type lamins, and experimentally test the impact of each on nuclear mechanics. An additional limitation of our study is that we express human lamins in mouse cells, and this approach might overlook subtle species-specific differences between lamins. Finally, all the exogenous lamin constructs possess an N-terminal FLAG tag, which is known to slightly impair the rescue ability of LaA^28^. In our system, the FLAG tag is necessary to assess the relative abundance of the exogenous lamins and to allow for direct comparison in the co-IP experiments. By using the same small tag for all expression constructs, we aimed to avoid any potential bias. Furthermore, since we previously showed that the FLAG-tag slightly reduces the ability of LaA to provide structural support to the nucleus^28^, we anticipate that the actual rescue effect of LaA on nuclear deformability is even higher than reported here.

In summary, nuclear deformability is important in several cellular contexts, such as protecting nuclei from external mechanical stress, allowing for nuclear deformation during migration through confined environments, and sensing external mechanical forces and confinement^29^. Here, we demonstrate that while A-type and B-type lamins jointly modulate the stiffness of the nuclear interior, A-type lamins are able to resist large strains to the nucleus in a regime dependent on the identity of the lamin rod domain. This may stem from different networks formed from lamins possessing the A-type or B-type rods that confer different mechanical strength to the nucleus, or via differences in the strength or assembly of the filaments themselves. Current electron tomography studies are unable to resolve structural differences between A-type and B-type lamins filaments in situ^6^, but future refinements might offer new insights into distinguishing features between their respective filaments and networks. Notably, both lamin types can properly position nuclear pores, while also possessing several unique interaction partners that likely require the presence of a tail domain. We envision a separation of function between the different lamin domains, with rods as key drivers for the ability of lamins to resist deformation, and the tails as hubs for protein-protein interactions. It will be of particular interest to examine LaA-specific interactors in future work, as these may provide clues to pathways and processes that are dysregulated in laminopathies and cannot be compensated for by B-type lamins.

## Extended Data Figures

**Extended Data Figure 1:**
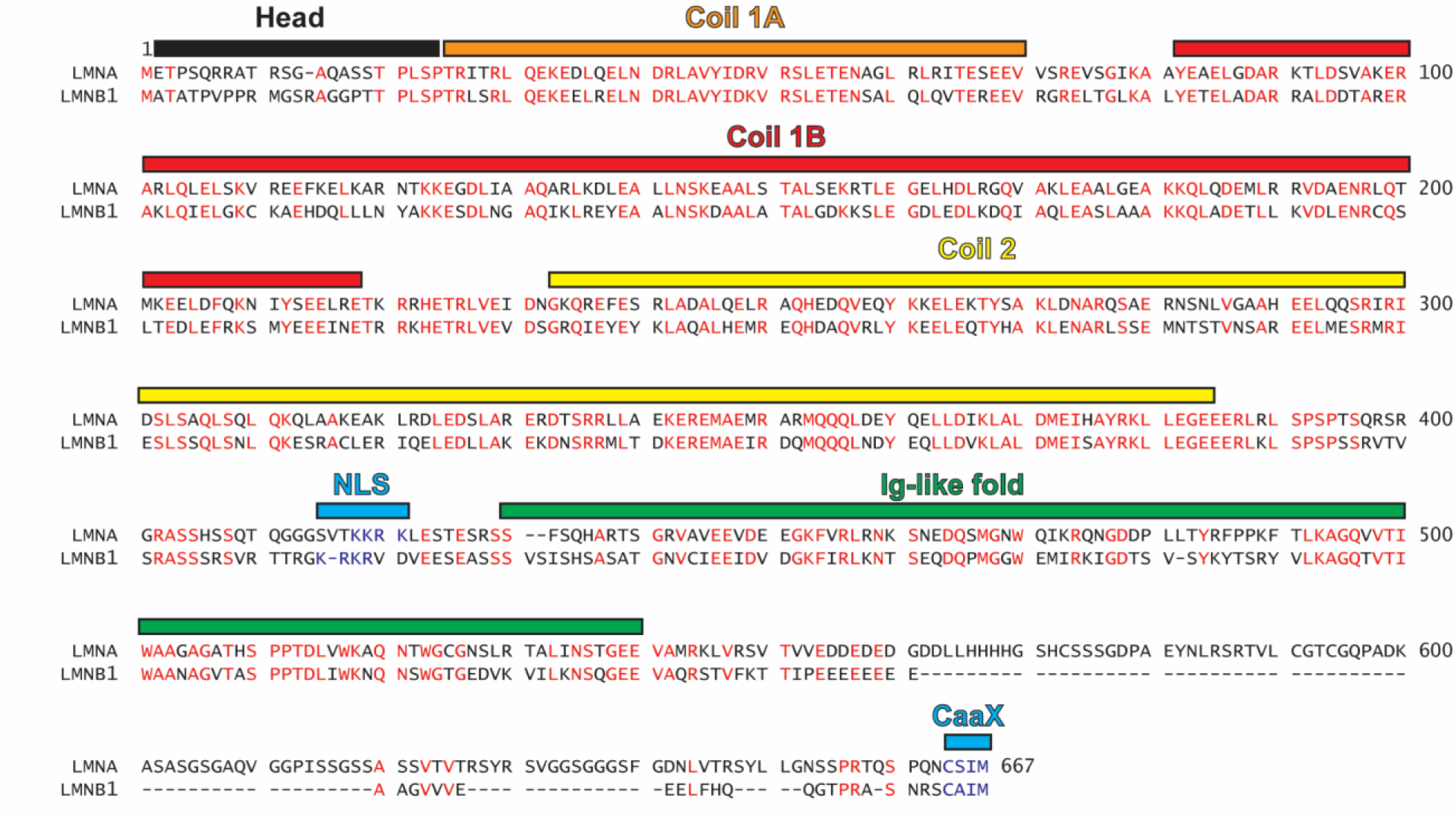
Sequence alignment between human LaA and LaB1. Amino acids colored in red are identical between the two proteins. The positions of relevant lamin domains are indicated by colored bars above the sequence. NLS = nuclear localization signal; Ig-like fold = Immunoglobulin-like fold.

**Extended Data Figure 2:**
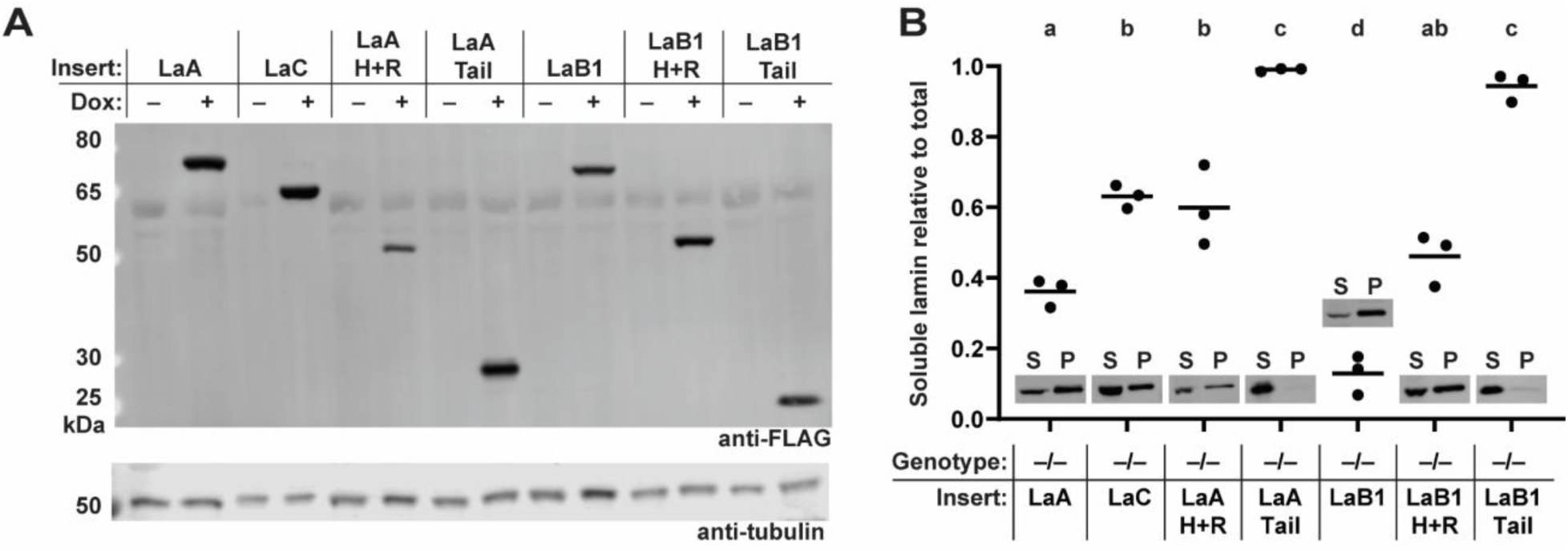
Lamin constructs are expressed at similar levels but differ in their solubility. (**A**) Lysates from *Lmna*^-/-^ MEFs expressing the indicated constructs were separated by SDS-PAGE and membranes were probed with anti-FLAG antibody to visualize the various lamin constructs. A ‘no dox’ control was included for each cell line, which indicates that there is no exogenous lamin expression in the absence of doxycycline in the media. Tubulin was used as loading control. (**B**) Results from differential protein extraction experiment using buffers of different stringency. The ‘easily soluble’ (S) fraction was collected from lysates using a buffer with 1% NP-40, which has been shown previously to solubilize the nucleoplasmic pool of lamins. The ‘insoluble fraction’ (P) was then resuspended in a high-salt RIPA buffer, and both fractions were separated by SDS-PAGE and probed with anti-FLAG antibodies to determine the relative abundance of each lamin in each fraction. Points on the graph represent the soluble lamin relative to the total signal observed in both fractions, from 3 independent experiments. Bars indicate overall fraction of soluble lamin, and insets show representative bands from one replicate. Sets of points with the same letter above them are not significantly different, whereas different letters indicate *p* < 0.05, One-way ANOVA with Tukey’s multiple comparison test.

**Extended Data Figure 3:**
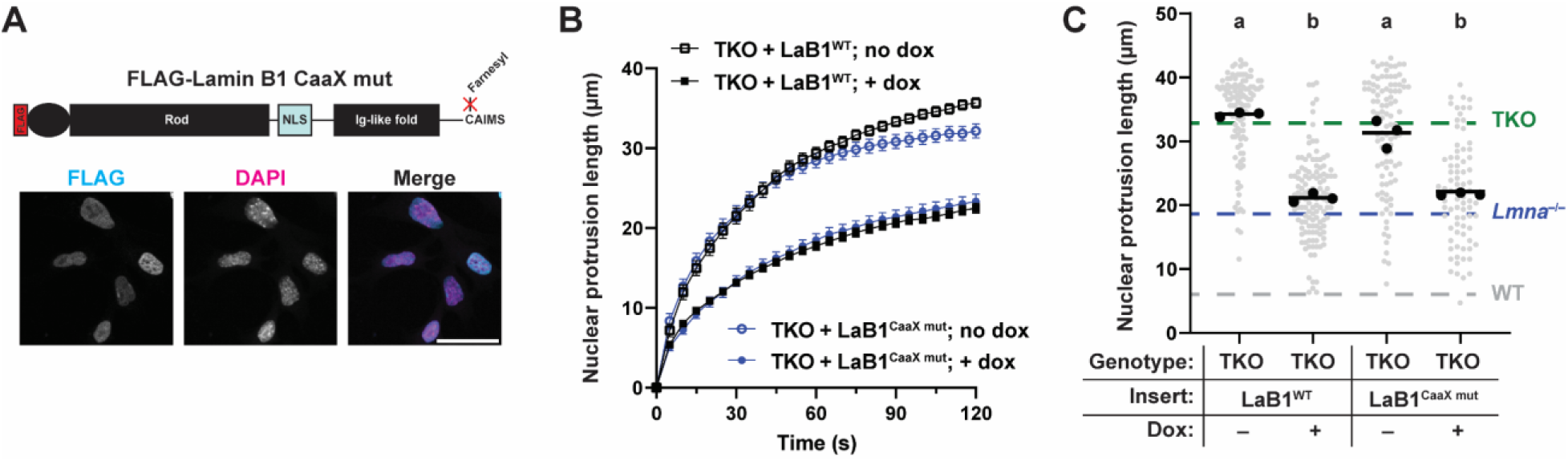
LaB1^CaaX^ ^mut^ exhibits abnormal localization yet rescues nuclear deformability similar to wild-type LaB1. (**A**) Schematic of LaB1^CaaX^ ^mut^ which contains an extra serine at the end of the CaaX motif (CAIM ➔ CAIMS). This mutation abolishes the normal nuclear rim staining of LaB1, as seen in the immunofluorescence image. Scale bar = 50 µm. (**B-C**) Quantification of nuclear deformability of TKO MEFs expressing LaB1^WT^ or LaB1^CaaX^ ^mut^. Grey points indicate measurements from individual cells, black points indicate replicate means, and bars indicate overall means. Sets of points with the same letter above them are not significantly different, whereas different letters indicate *p* < 0.05, One-way ANOVA with Tukey’s multiple comparison test.

**Extended Data Figure 4:**
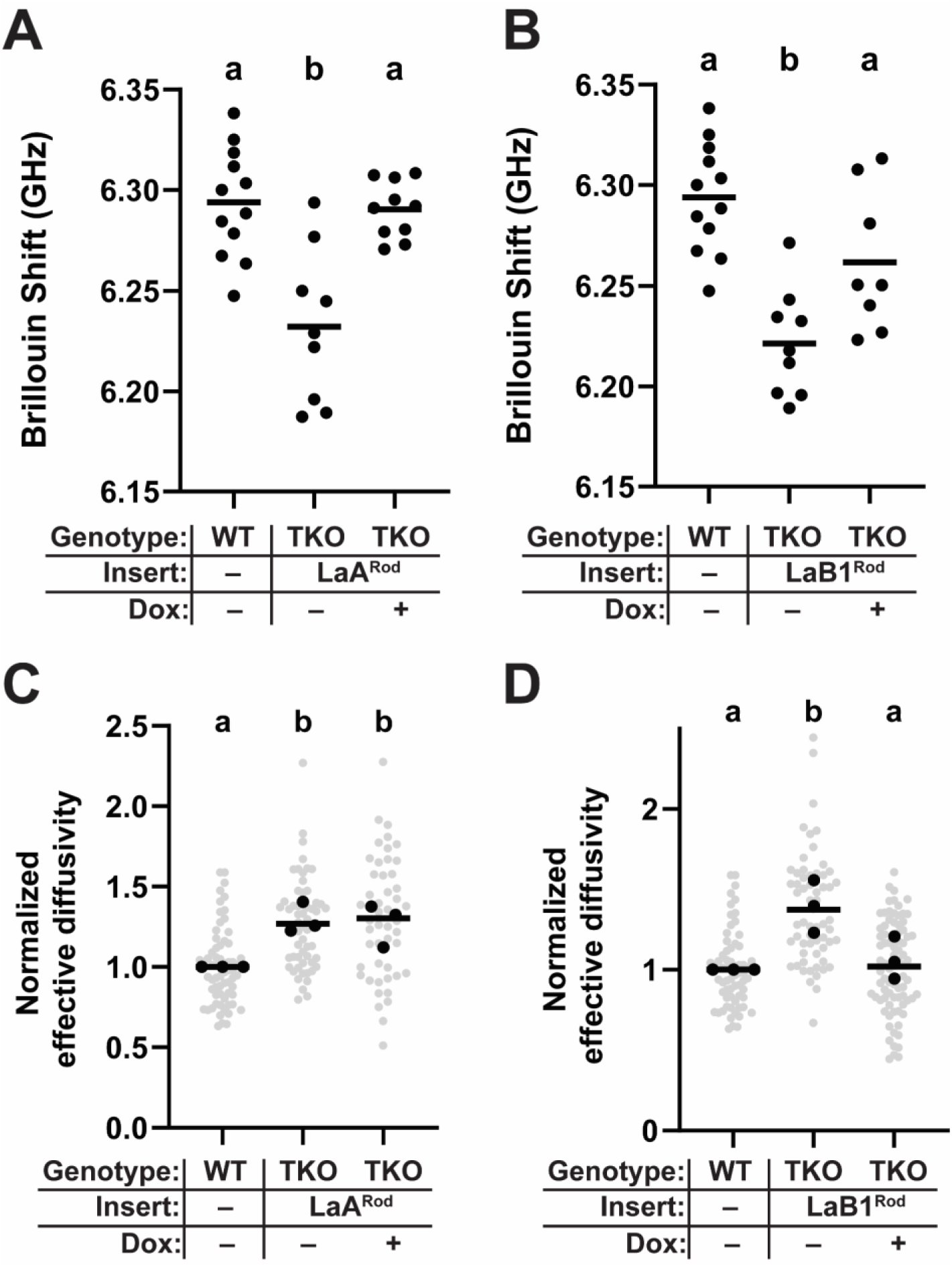
Brillouin and nucGEMs analysis results for lamin Head+Rod truncations. (**A-B**) Brillouin frequency shifts within the nucleus of TKO MEFs expressing LaA^Head+Rod^ (**A**) or LaB1^Head+Rod^ (**B**), as in Figure 2. Points represent measurements from individual cells, bars indicate means, and different letters above the sets of points indicate *p* < 0.05 One-way ANOVA with Tukey’s multiple comparison test. (**C-D**) Quantification of normalized nucGEM effective diffusivity of TKO MEFs expressing LaA^Head+Rod^ (**C**) or LaB1^Head+Rod^ (**D**), as in Figure 2. Grey points indicate measurements from individual cells, black points indicate replicate means, and bars indicate overall means.

**Extended Data Figure 5:**
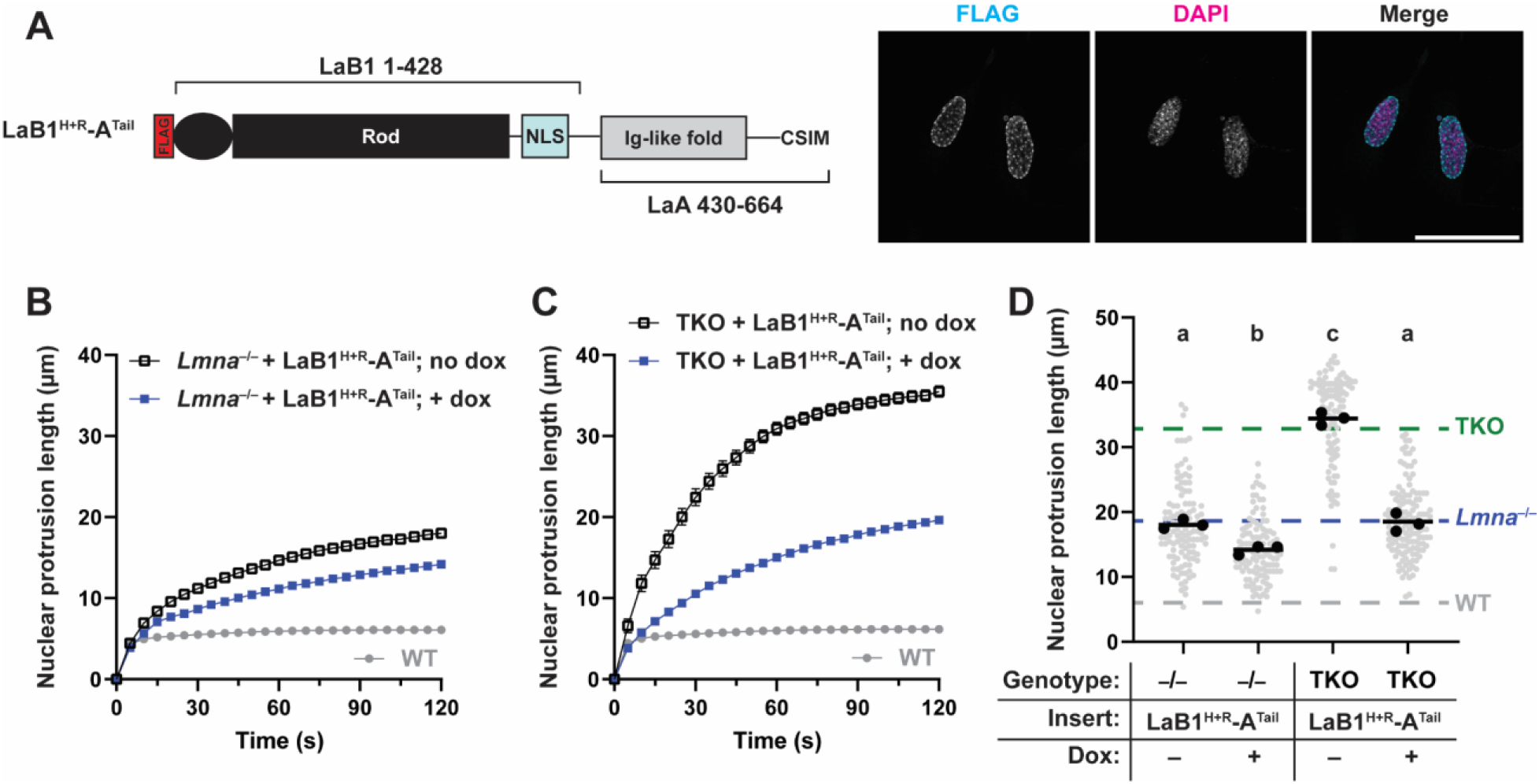
LaB1^Head+Rod^-LaA^Tail^ chimera most closely resembles LaB1. (**A**) Schematic depiction of LaB1^Head+Rod^-LaA^Tail^ chimera and representative image of *Lmna*^-/-^ MEFs expressing this construct. Scale bar = 50 µm. Note the punctate localization of this chimera, which resembles the LaB1^Head^-LaA^Rod+Tail^ construct. (**B-D**) Quantification of nuclear deformability of *Lmna*^-/-^ (**B**) or TKO (**C**) MEFs expressing LaB1^Head+Rod^-LaA^Tail^ chimera. (**D**) Quantification of nuclear protrusion lengths of all cells 100 s after the start of aspiration. The rescue achievable by this chimera is most similar to LaB1. Grey points indicate measurements from individual cells, black points indicate replicate means, and bars indicate overall means. Sets of points with the same letter above them are not significantly different, whereas different letters indicate *p* < 0.05, One-way ANOVA with Tukey’s multiple comparison test.

**Extended Data Figure 6:**
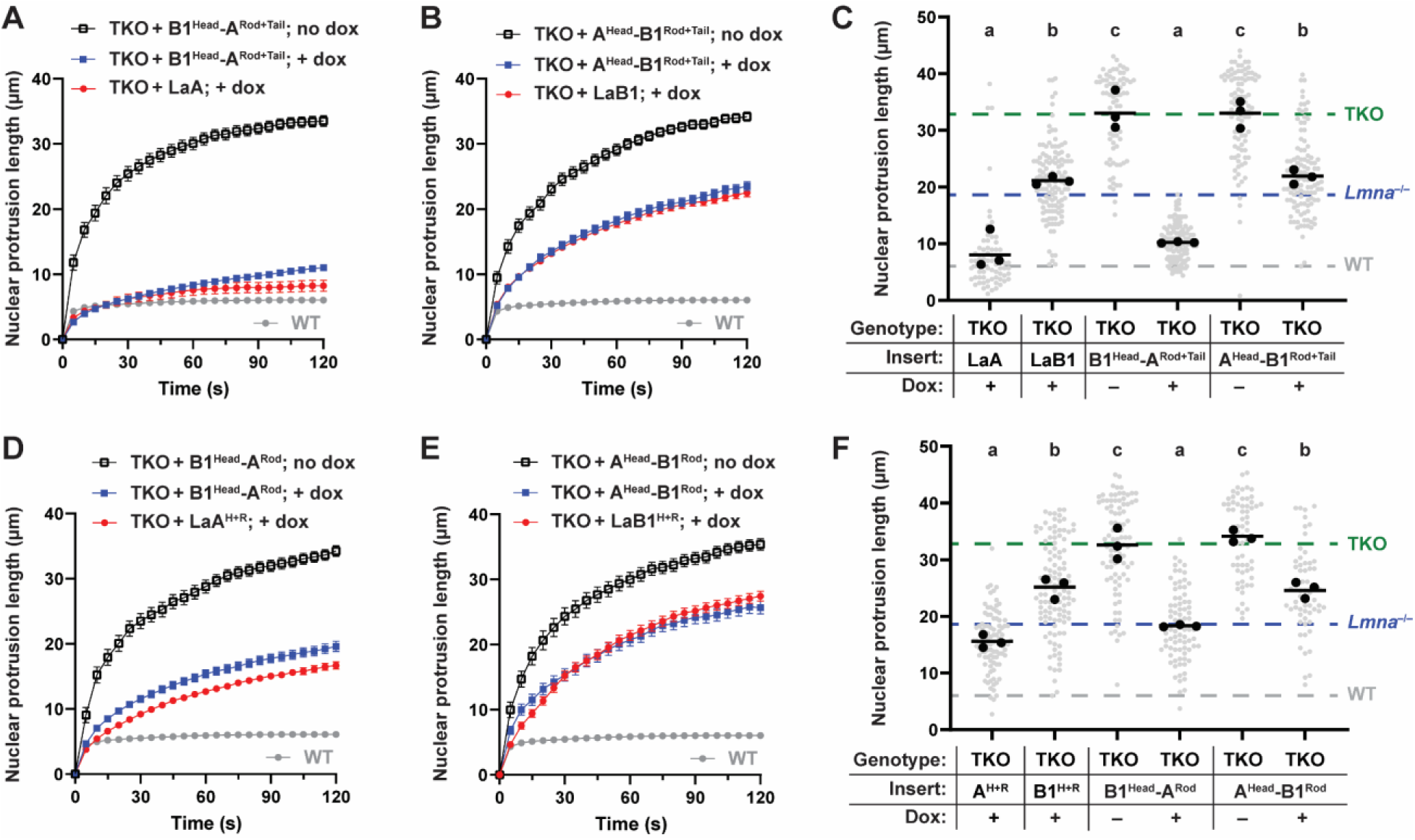
Nuclear deformability of TKO MEFs expressing head-swapped lamins. (**A-C**) Time-dependent nuclear protrusion of TKO MEFs expressing LaB1^Head^–LaA^Rod+Tail^ (**A**) or LaA^Head^–LaB1^Rod+Tail^ (**B**), with quantification of nuclear protrusion length following 100 s of aspiration (**C**). Data from TKO MEFs expressing LaA or LaB1 are shown for reference. (**D-F**) Time-dependent nuclear protrusion of TKO MEFs expressing LaB1^Head^ – LaA^Rod^ (**D**) or LaA^Head^ –LaB1^Rod^ (**E**), with quantification of nuclear protrusion length following 100 s of aspiration (**F**). Data from TKO MEFs expressing LaA^Head+Rod^ or LaB1^Head+Rod^ are shown for reference. Grey points indicate measurements from individual cells, black points indicate replicate means, and bars indicate overall means. Sets of points with the same letter above them are not significantly different, whereas different letters indicate *p* < 0.05, One-way ANOVA with Tukey’s multiple comparison test.

**Extended Data Figure 7:**
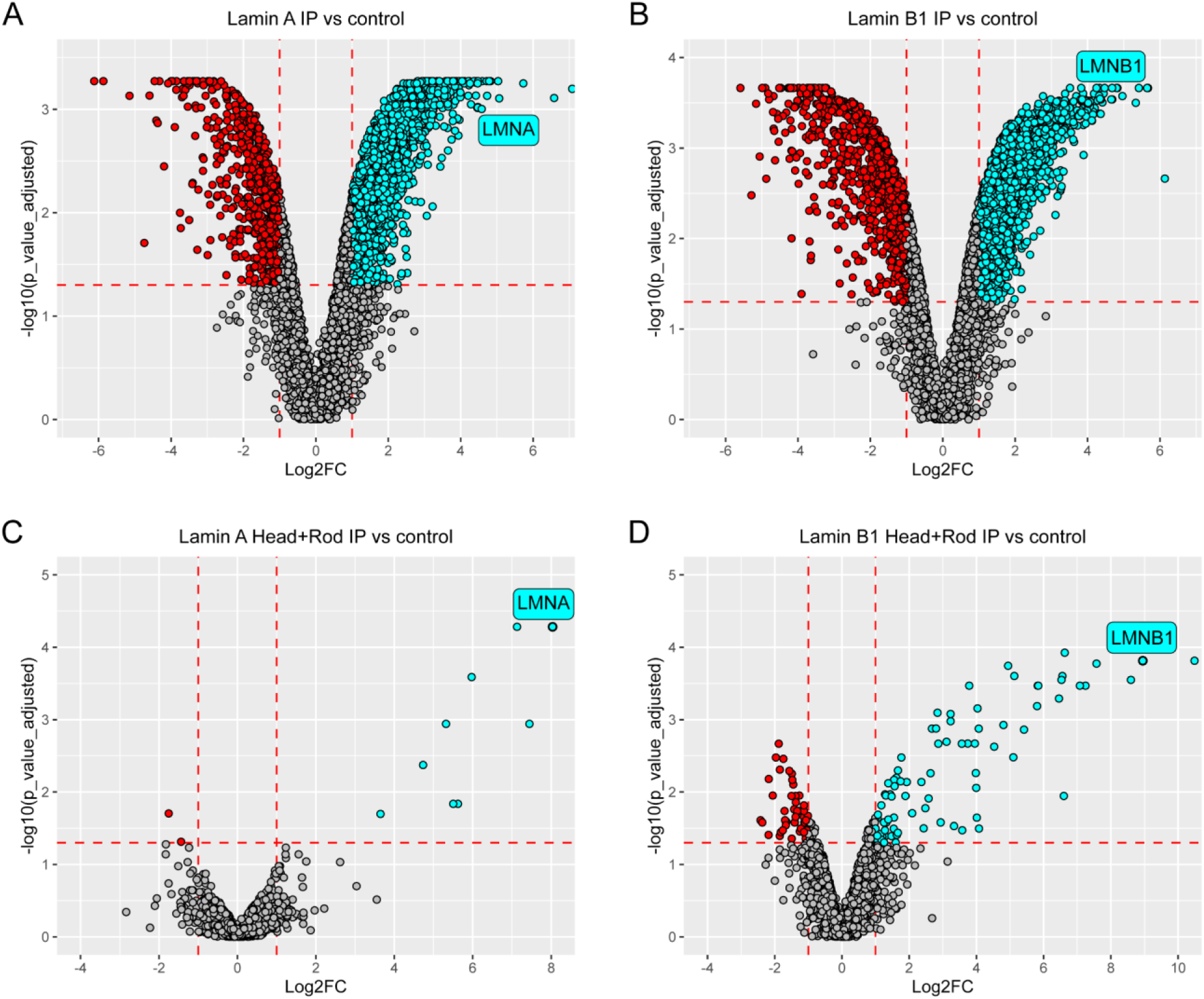
Volcano plots comparing lamin co-IP’s to negative control co-IP. Negative control co-IP’s were performed using the same anti-FLAG beads using lysates from parental *Lmna^-/-^* MEFs that did not express any FLAG-tagged proteins. Red points were significantly enriched in the negative control and cyan points were proteins enriched in each co-IP. The bait proteins are indicated for LaA (**A**), LaB1 (**B**), LaA^Head+Rod^ (**C**) and LaB1^Head+Rod^ (**D**). Dashed red lines indicate the fold change and significance thresholds to identify hits.

**Extended Data Figure 8:**
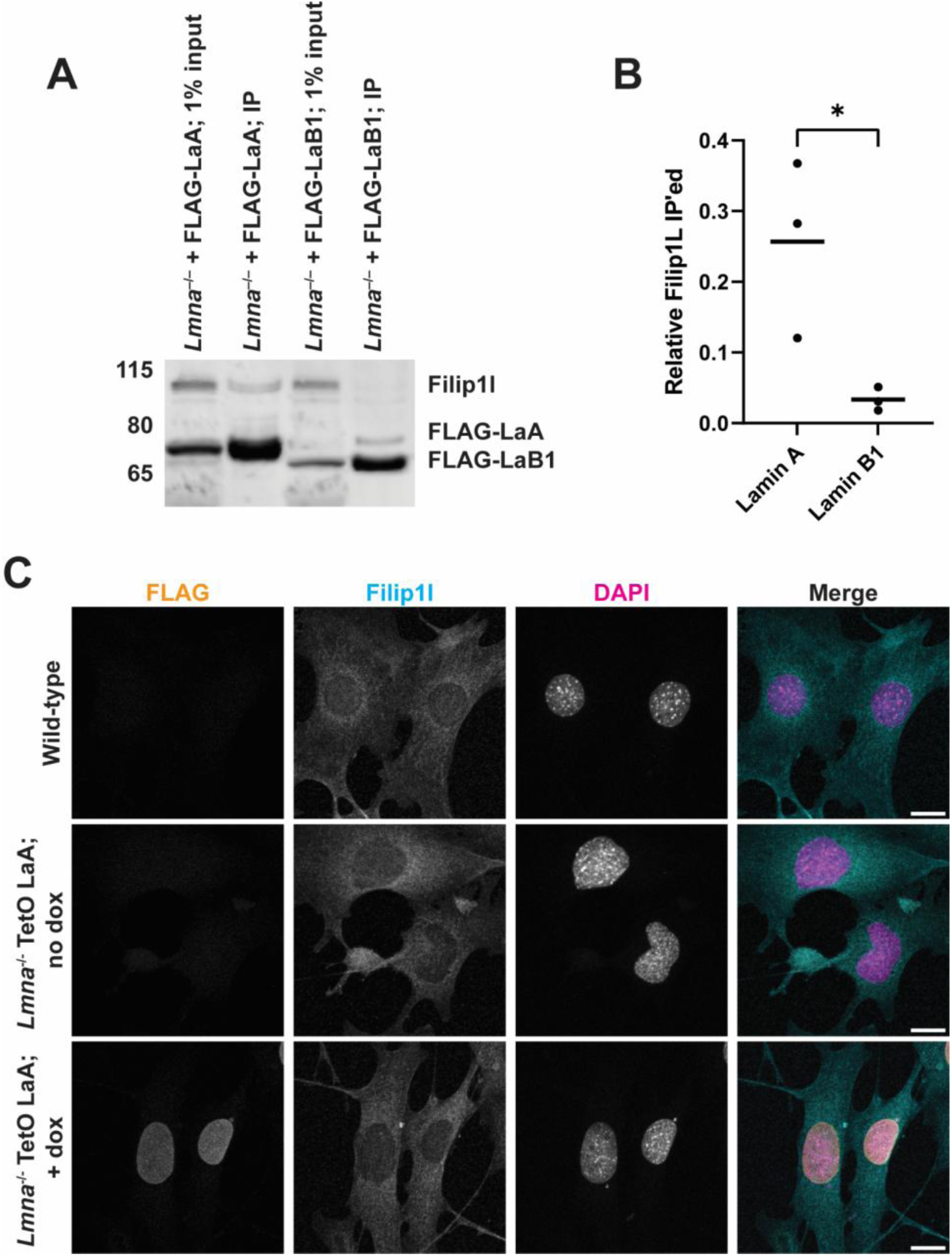
Filamin A interacting protein 1-like (Filip1l) preferentially interacts with A-type lamins. (**A**) Representative immunoblot of co-IP experiment performed using FLAG-LaA or FLAG-LaB1 as the bait protein. (**B**) Quantification of Filip1l immunoprecipitated by LaA versus LaB1. Mean intensity of the Filip1l band was normalized (divided) by the mean intensity of the bait protein in the IP eluate. Each point represents data from independently performed co-IPs and immunoblots. (**C**) Immunofluorescence staining for Filip1l in wild-type, *Lmna*^-/-^ MEFs, and *Lmna*^-/-^ MEFs expressing LaA. Filip1l had a cytoplasmic localization in each cell line, consistent with previous reports ^87^, with no obvious differences in localization following loss of Lamin A/C or reintroduction of LaA. Scale bar = 10 µm.

## Methods

### Cell culture

Wild-type and *Lmna*^-/-^ MEFs were a kind gift of Colin Stewart ^17^. TKO MEFs were a kind gift of Stephen Young and Loren Fong ^36^. Cells were maintained in DMEM supplemented with 10% FBS and 1% penicillin/streptomycin in a humidified incubator set to 37°C. Cells were passaged at 80-90% confluency and routinely checked for mycoplasma contamination by PCR. For stable genetic manipulations, the PiggyBac transposase system was used as described below. Antibiotic selection was performed using Puromycin at 3 µg/ml for at least 1 week, or until cells in a non-transformed well had all died. To induce expression of the exogenous lamin, doxycycline was added at 100 ng/ml for 24 h prior to experiments. To block farnesylation, Lonafarnib (Sigma SML1457) was dissolved in DMSO and added at a final concentration of 25 µM to cell culture media. Lonafarnib was added simultaneously with doxycycline, so that all the newly synthesized lamin would not be farnesylated.

### Genetic construct information

The doxycycline inducible FLAG-LaA and LaB1 constructs have been described previously^42^, and these constructs served as the basis for PCR amplification to clone lamin truncations and chimeras. All constructs were cloned via Gibson assembly into the pPB-rtTA-hCas9-puro-PB backbone^88^ following digestion with NheI and AgeI. A list of primers used for amplification of the various constructs is presented in Supplemental Table 1. Following cloning, all constructs were verified to have to correct sequence by Sanger sequencing of the inserts. For PiggyBac transposition, plasmids containing the desired insert were co-transfected into cells with a plasmid encoding a hyperactive transposase (2:1 vector plasmid: hyperactive transposase plasmid) using Mirus LT1 transfection reagent according to manufacturer’s instructions.

### Micropipette aspiration

Micropipette aspiration was performed using a previously published protocol^38^. Briefly, approximately 1 × 10^6^ cells were suspended in buffer containing 2% BSA in PBS supplemented with 0.2% FBS and 10 mM EDTA to prevent cell adherence. Hoechst 33342 was added to the cells immediately before the cell suspension was transferred to the micropipette device. Cells were perfused using the following pressure settings, controlled by a Fluigent microfluidics controller: top port: 1.0 psi, bottom port: 0.2 psi. This pressure gradient drives the perfusion of cells into the micropipette ‘pockets’ that contain the aspiration channels. Once a flow of cells was established in the device, cells were cleared from the pockets to allow new cells to enter, and images of nuclear protrusion over time were acquired every 5 s for 40 frames. Nuclear protrusion length was measured using a MatLab script available at (https://github.com/Lammerding/MATLAB-micropipette_analysis).

### Immunofluorescence analysis

Cells were seeded on fibronectin-coated 12 mm glass coverslips overnight, then doxycycline (100 ng/ml) was added for 24 hours to induce expression of the exogenous lamin. Cells were fixed in 4% paraformaldehyde in PBS for 15 min at room temperature, followed by three 5-min washes with IF wash buffer containing 0.2% Triton X-100, 0.25% Tween 20 and 0.3% BSA in PBS. Cells were blocked in 3% BSA in PBS for 1 h, then primary antibodies were added for 1 h in blocking buffer at room temperature or overnight at 4°C. Primary antibodies used were: anti-FLAG (Sigma F7425, 1:1000), anti-NPC (Abcam AB24609, 1:1000), anti-FILIP1L (Proteintech 30134-1 AP, 1:250). DAPI was added 1:500 in PBS for 15 minutes to stain DNA. Secondary antibodies used were Alexa Fluor 488 or 568-conjugated donkey anti mouse or rabbit-IgG antibodies (Invitrogen) diluted 1:250 in 3% BSA in PBS. Coverslips were mounted on glass slides using Mowiol and kept in the dark until imaging.

### Co-immunoprecipitation

Cells (5 × 10^5^) were seeded in 10-cm plates, allowed to adhere overnight, then doxycycline was added (100 ng/ml) for 24 hours to induce expression of FLAG tagged lamin. Cells were lysed on ice in high-salt RIPA buffer containing 12 mM sodium deoxycholate, 50 mM Tris-HCl pH 8.0, 750 mM NaCl, 1% (v/v) NP-40 Alternative, 0.1% (v/v) SDS. Lysates were then vortexed for 5 min, sonicated (Branson 450 Digital Sonifier) for 30 s at 36% amplitude, and centrifuged at 4°C for 10 min at 14,000 g. For each co-IP, 20 µL of anti-FLAG beads (Sigma M8823) were washed twice in high-salt RIPA buffer (300 µL) to equilibrate the beads. 1% of the whole cell lysate was stored at -80°C, and the remainder was mixed with the equilibrated anti-FLAG beads and the total volume was brought up to 1 ml. Immune complexes were allowed to form overnight at 4°C with gentle agitation. Proteins bound to the beads were then washed 5 × with co-IP wash buffer (50 mM Tris-HCl pH 8.0, 300 mM NaCl, 0.3% (v/v) Triton X). Bound proteins were eluted from beads using 100 µL low-pH elution buffer (0.1 M Glycine pH 3.0) for 10 minutes with orbital rotation at 40 rpm. Eluates were immediately quenched with 20 µL 1M Tris (pH 8.0) to neutralize the samples.

For mass spectrometry identification of interactors, protein concentration was measured using BCA assay, and 3 µg of protein for each sample were added to a final volume of 100 µL. 8M urea in 100 mM Tris (pH 8.0) was added to a final concentration of 2M. TCEP (Thermo Scientific, Cat# 77720) was added to a final concentration of 5 mM, and the sample was incubated at room temperature for 30 minutes. Iodoacetamide (Cytiva, Cat# RPN6302) was then added to a final concentration of 10 mM, followed by incubation at room temperature for 30 minutes in the dark. Digestion was performed by adding 1 µL of Trypsin Gold (Promega, Cat# V5280, 1 µg/µL), and the sample was incubated overnight at 37°C. Finally, an equal volume of 4% formic acid solution was added, and the sample was mixed and stored at -80°C until mass spectrometry analysis.

### Mass Spectrometry Analysis

Peptides were desalted using a homemade spin column, resuspended in 10 µL of Solution A (0.1% formic acid in water), and 3 µL (around 500 ng peptides) was injected for mass spectrometry analysis. The digested peptides were analyzed using nanoLC-ESI-MS/MS on a timsTOF HT (Bruker) coupled with a nanoElute2 ultra-high-performance liquid chromatography system or Evosep One chromatography system. For nanoElute2 ultra-high-performance liquid chromatography system, separation was performed using a homemade C18 column (100 μm ID, 10 cm length, 1.9 μm C18 particle size) at a flow rate of 500 nL/min with mobile phases of Solution A (0.1% formic acid in water) and Solution B (0.1% formic acid in acetonitrile). The gradient elution was as follows: 2% to 35% B over 25 min, 35% to 95% B over 0.5 min, followed by a 4.5 min hold at 95% B. Samples were first loaded onto a Thermo PepMap™ Neo Trap Cartridge with PepMap™ Neo UHPLC Columns (Thermo Scientific, Cat# 174500) before separation on the homemade analytical column.

For Evosep One chromatography system, EvoTips were conditioned with 100% isopropanol for 1 min, then washed twice with 50 μL of buffer B (Mass Spec Grade Acetonitrile with 0.1% formic acid) by centrifugation at 700g for 60 s. The washed EvoTips were equilibrated by washing three times with 50 μL of buffer A (Mass Spec Grade Water with 0.1% formic acid), followed by centrifugation at 700g for 60 s each time. Next, around 500 ng peptides were loaded onto the EvoTips and centrifuged at 700g for 60 s. The loaded peptides were washed twice with buffer A by centrifugation at 700g for 60 s each. To keep the peptides wet, 200 μL of buffer A was applied to the top of the EvoTip and centrifuged at 700g for 30 s. The peptides on the EvoTips were separated using a homemade 8 cm × 150 μm analytical column packed with 1.5 μm C18 beads. Separation was performed over 22 min following the manufacturer’s standard 60SPD method. The analytical column was equilibrated at 2 μL/min, with a gradient flow of 1 μL/min, which was increased to 2 μL/min for washing. Peptides were eluted using solvent A (Mass Spec Grade Water with 0.1% formic acid) and a gradually increasing concentration of solvent B (Mass Spec Grade Acetonitrile with 0.1% formic acid).

### DIA Data Acquisition and analysis

DIA data acquisition on the timsTOF HT was performed using Compass Hystar version 6.3 (Bruker) and timsControl version 6.0.2 (Bruker). The DIA-PASEF method covered an m/z range of 100–1700, with 60 windows spanning 300–1300 Da and an ion mobility range (1/K₀) of 0.60–1.45 V·s/cm². The ramp time was 75.0 ms, and the accumulation time was 50.0 ms. The capillary voltage of the CaptiveSpray source was 1500 V, with a dry gas temperature of 180°C and a flow rate of 3.0 L/min. The collision energy was set to 10.0 eV. Before each set of runs, m/z and mobility calibration was performed using three reference ions from the ESI-Filter Cal (m/z 622, 922, 1222). DIA data was analyzed using DIA-NN (version 1.8.1) with a Bruker mouse proteome spectral library (Bruker_Mouse_Trypsin_TIMScore_v2.tsv). The parameters included up to 3 missed cleavages, a maximum of 2 variable modifications, a precursor charge range of 1–4, and a precursor m/z range of 100–1800. The analysis was performed in double-pass mode for the neural network classifier, with ‘Any LC (high accuracy)’ selected for quantification. Match Between Runs (MBR) was enabled. All other settings were kept at default, with RT-dependent cross-run normalization and results filtered at 1% FDR. The analysis was executed using 32 threads, as automatically suggested by the software.

### Statistical Analysis of mass spectrometry results

We employed a linear model-based pipeline tailored to our experimental design to analyze protein quantification results obtained from DIA-NN. Protein groups passing a 1% false discovery rate (FDR) threshold were retained for downstream analysis and further filtered based on two criteria: (1) the number of valid protein intensity values per run and (2) the median protein intensity per run. Proteins with fewer than three valid measurements across the bait replicates were excluded from subsequent steps. For fold change (FC) estimation, protein intensity ratios were computed using a specific one-to-one replicate pairing strategy. For example, in the case of three replicates for both the Wild-Type (WT) and Control (C) conditions, ratios were calculated as WT1/C1, WT2/C2, and WT3/C3. The final fold change for each protein was defined as the median of all computed replicate ratios. To assess statistical significance, a two-sample t-test was performed by comparing the distribution of each protein’s ratios to those of all other proteins in the dataset, yielding raw p-values. Statistical significance was further refined using empirical Bayes moderation of the variance, as implemented in the *limma* package (v3.62.2) within the R environment (v4.4.2).

To quantify differences in protein abundance between two distinct bait proteins, we used the previously described pipeline to calculate FCs and FDRs. Protein interactors for each bait were identified by comparison to control samples, applying thresholds of FC > 2 and FDR < 0.05. The analysis then focused on the union of interactors identified under both bait conditions. To assess differences in protein interaction profiles between the two baits, we directly compared their respective runs. Data were normalized such that the median protein intensity was approximately equal across all runs, ensuring comparability between conditions. Results were visualized using a volcano plot, with fold changes plotted against adjusted p-values. Known AP-MS contaminants— such as keratins (KRT), small ribosomal subunit proteins (RPS), and large ribosomal subunit proteins (RPL)—were excluded from the final dataset. Comparison of interactors identified using each lamin bait was performed using InteractiVenn^89^.

### Immunoblotting and differential protein extraction

Cells (10^5^) were seeded in wells of a six-well plate, and doxycycline was added at the appropriate concentration for 24 h to induce protein expression. To isolate the easily soluble fraction, cells were lysed using 200 μl low-salt buffer (0.5×PBS, 50 mM HEPES pH 8.0, 10 mM MgCl2, 1 mM EGTA and 0.2% NP-40 Alternative). Cells were lysed on ice for 5 min, then cells were scraped off the plate, transferred to 1.7 ml microcentrifuge tubes, and spun at 4°C for 5 min at maximum speed (14,000 g for 5 min) in a benchtop centrifuge. The supernatant was saved as the ‘soluble fraction’. The pellet was resuspended in 200 μl high-salt RIPA buffer [12 mM sodium deoxycholate, 50 mM Tris-HCl pH 8.0, 750 mM NaCl, 1% (v/v) NP-40 Alternative, 0.1% (v/v) SDS], vortexed for 5 min, sonicated (Branson 450 Digital Sonifier) for 30 s at 36% amplitude, boiled for 2 min, and centrifuged at 4°C for 10 min at 14,000 g. The supernatant from this step was saved as the ‘insoluble fraction’. Equal amounts of each fraction (20 μl) were mixed with 5×Laemmli buffer, boiled for 3 min and then separated by SDS-PAGE. Samples were denatured by boiling for 3 min, loaded onto 4–12% Bis-Tris gels (Invitrogen NP0322), run for 1.5 h at 100 V, then transferred for 1 h at 16 V onto PVDF membrane. Membranes were blocked for 1 h in blocking buffer containing 3% BSA in Tris-buffered saline plus 1% Tween 20. Primary antibodies used: anti-FLAG (Sigma F7425, 1:2000), anti-tubulin (Sigma T6199, 1:3000). Secondary antibodies used were: Licor IRDye 680RD donkey anti-mouse-IgG (926-68072; 1:5000) and Licor IRDye 800CW Donkey anti-Rabbit IgG (926-32213; 1:5000). Secondary antibodies were added for 1 h at room temperature in blocking buffer, followed by three 10-min washes. Membranes were imaged using the Odyssey Licor scanner, and then cropped and brightness and contrast was adjusted using Image Studio Lite (version 5.2) software.

### Brillouin Microscopy

Spontaneous Brillouin light scattering arises from the interaction between incident light and intrinsic acoustic phonons within a sample. This interaction leads to a Doppler-like GHz range frequency shift in the scattered light, known as the Brillouin frequency shift (𝑣_*B*_) proportional to the longitudinal elastic modulus (*M*^′^) and is given by 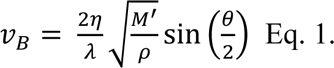 where *η* and *ρ* are refractive index and density of material respectively, *λ* is laser wavelength, and *θ* is the collection angle of scattered light (180°).

A home-built confocal Brillouin microscope was used for acquiring the Brillouin frequency shift of wild-type and TKO cells expressing full length or truncated lamins. During imaging, the cells were in stage-top Okolab Incubator at 37°C temperature and 5% CO_2_. The experimental setup is shown in Supplemental Figure 2. The input light source for Brillouin microscope was 660 nm laser (Torus, Laser Quantum Inc.) with a power of ∼20 mW on sample. The laser beam was focused on sample by an objective lens (Olympus, 40x/0.95 NA) mounted on an inverted microscope (Olympus, IX81), achieving a focal spot size of approximately 0.42 µm × 0.42 µm × 1.46 µm. The backscattered Brillouin signal was collected by the same objective analyzed by two-stage virtually imaged phase array-based spectrometer with 15 GHz of free spectral range. The Brillouin spectrum is recorded by an electron-multiplying charge coupled device camera (iXon Andor) with an exposure time of 0.05 s. The microscope was calibrated with electro-optic modulator (EOM) before each measurement. Two-dimensional Brillouin images of cells were acquired by using XY motorized stage with a step size of 1-2 µm. A brightfield microscope, co-aligned with the Brillouin scanning arm, and a CMOS camera (Andor Neo) were used to acquire structural images. To visualize the nucleus, cells were stained with the fluorescent dye (Hoechst 33342, Thermo Fisher Scientific) 30 minutes prior to imaging. Fluorescence images were acquired using the same optical path by switching from the brightfield to the fluorescence channel by rotating the filter turret 2 as shown in Supplemental Figure 4.2.

### Brillouin data acquisition and analysis

LabVIEW-based acquisition program (National Instruments, version 2024) was developed in-house to capture brightfield images, fluorescence images, and Brillouin spectra. Spectrometer was calibrated using EOM allowing determination of the free spectral range and the pixel-to-frequency conversion factor. Brillouin shifts at each pixel were extracted by fitting the acquired spectra to a Lorentzian function using MATLAB (MathWorks, R2024 a). The resulting pixel-wise Brillouin shift values were used to reconstruct two-dimensional Brillouin images. Co-registration of Brillouin images with corresponding brightfield and fluorescence images was conducted in MATLAB. Brightfield and fluorescence images were resized to match the spatial resolution of the Brillouin images, aligned accordingly, and overlaid to delineate subcellular regions. This co-registration enabled precise extraction and quantification of Brillouin shifts of cytoplasm and nucleus of cells. The average Brillouin frequency shift of cytoplasm, nucleus, and entire cell was calculated for all cells and used to quantify their mechanical properties.

### NucGEMs experiments and analysis

*Quasibacillus thermotolerans* (QtE) nuclear Genetically Encoded Multimeric nanoparticles (nucGEMs)^54^ were introduced into MEFs via reverse transduction using a Lentiviral plasmid delivery system. After cell lines stabilized, they were either used for experiment or frozen in 10% DMSO in FBS for future use and thawed for use in experiments as needed. For QtE-nucGEM analysis, approximately 15-20k cells were seeded in a 24-well glass bottom plate, and the next day, 100 ng/ml doxycycline was added to induce the expression of lamin constructs. After 24 h, cells were moved to the confocal microscope unit fitted with incubators for image acquisition. Cells were maintained at 37°C and 5% CO_2_ incubators throughout and also during the entire period of image acquisition. Micrographs were acquired on a Nikon Eclipse Ti Eclipse microscope mounted with Yokogawa CSU-X1 spinning disk unit, NIDAQ AOTF multilaser unit, and Prime 95B camera operating on Nikon NIS-Elements AR (v 5.21.03) software. We used CFI Apo 60x/NA-1.49/.12 TIRF objective with a 470/40m excitation filter and ET525/36m emission filter (Chroma Technology Corp) in all mammalian acquisitions. Using a 488 nm laser the sapphire fluorophore was excited using 100% power and images were collected from a single focal plane at 100fps, binning 1, 512×512, and 8-bit pixel depth for 2 seconds. To analyze GEM movement, GEMs were initially tracked with the ImageJ (2.1.0/1.54j Particle Tracker 2D-3D tracking algorithm from MosaicSuite. Nuclear staining was used to segment the tracks within the nuclear region. Trajectories were then analyzed with the GEM-Spa (GEM single particle analysis) software package that we developed in house: https://github.com/liamholtlab/GEMspa

### Microscopy

Confocal immunostaining images were acquired on a Zeiss LSM900 series confocal microscope with airyscan module using a 40× water immersion objective. The optimal z-slice size was automatically determined using Zen Blue (Zeiss) software. Airy units for images were set between 1.5 and 2.5. Micropipette aspiration data was acquired using an inverted Zeiss Observer Z1 epifluorescence microscope with Hamamatsu Orca Flash 4.0 camera. The image acquisition for micropipette aspiration experiments was automated with Zen Blue (Zeiss) software.

### Image analysis

Intensity profile measurements were performed using a FIJI macro available on request. Briefly, this macro used the ‘Plot Profile’ feature in FIJI software to measure the LaA intensity across a line drawn across a z-slice through the center of the nucleus. To account for differences in nuclear size, the intensity profiles are converted into relative nuclear distances by measuring the average intensity in each of 50 equally sized bins.

Qualitative assessment of NPC mislocalization was performed by observers blinded to experimental conditions. Nuclei were scored as having ‘normal’ or ‘mislocalized’ NPC distribution based on staining that was uniform along the nuclear periphery (normal), or clustered NPCs to one side of the nucleus and a clear lack of signal on the other side (mislocalized).

### Statistical analysis and figure generation

All analyses were performed using GraphPad Prism. Information on statistical tests used, cell counts, and significance values are present in each figure caption. Experiments were performed a minimum of three independent times, and for qualitative image analysis, observers were blinded to genotype and treatment conditions when scoring phenotypes. Our statistical analysis was developed in close consultation with the Cornell Statistical Consulting Unit. Figures were assembled using Adobe Illustrator.

## Acknowledgements

We thank Harald Herrmann and Ohad Medalia for helpful discussions regarding the structure of lamins. We thank the Biotechnology Resource Center (BRC) Flow Cytometry Facility (RRID: SCR_021740) and sequencing facility (RRID: SCR_021727) at the Cornell Institute of Biotechnology for their resources and technical assistance. This work was performed in part at the Cornell NanoScale Science & Technology Facility, a member of the National Nanotechnology Coordinated Infrastructure, which is supported by the National Science Foundation (award NNCI-2025233). This work was supported by awards from the Volkswagen Foundation (A130142 to J.L.), the National Institutes of Health (R01 AR084664, R01 HL082792, R01 GM137605, and R35 GM153257 to J.L., R01 GM132447 and R37 CA240765 to L.J.H, and R01 GM157084 to G.S.), the National Science Foundation (URoL2022048 to J.L.), the Maryland Stem Cell Research Fund (Discovery Grant to G.S.) and the Leducq Foundation (20CVD01 and 24CVD03 to J.L.). The content of this manuscript is solely the responsibility of the authors and does not necessarily represent the official views of the National Institutes of Health.

