## Supplementary Materials for "The Deformability of the Mammalian Cell Nucleus is Determined by the Identity of the Lamin Rod Domain"

#### Supplemental Table

| Primer Name | Sequence 5'-3 | Use |
|---|---|---|
| **LaA_fwd** | accctcgtaaaggtctagagaccatggactacaaagacgatg | Cloning LaC |
| **LaC_rev** | ccgtttaaactcattactaatcagcggcggctaccactcacgtggtggtgatggagcag | Cloning LaC |
| **LaA_rod_fwd** | accctcgtaaaggtctagagaccatggactacaaagacgatgacg | Cloning LaA H+R |
| **LaA_rod_rev** | ccgtttaaactcattactaattagctgctgcggctctcagt | Cloning LaA H+R |
| **LaA_IgFold_fwd** | accctcgtaaaggtctagagaccatggactacaaagacgatgacgacaagtctcactc atcccagacac | Cloning LaATail |
| **LaA_rev** | ccgtttaaactcattactaattacatgatgctgcagttc | Cloning LaATail |
| **LaB1_Rod_fwd** | accctcgtaaaggtctagagaccatggactacaaagac | Cloning LaB1 H+R |
| **LaB1_Rod_rev** | ccgtttaaactcattactaactaactactcgcctctgattc | Cloning LaB1 H+R |
| **LaB1_Tail_fwd** | accctcgtaaaggtctagagaccatggactacaaagacgatgacgacaagtcaagtc gtagtgtacgtac | Cloning LaB1Tail |
| **LaB1_Tail_rev** | ccgtttaaactcattactaactacataattgcacagcttc | Cloning LaB1Tail |
| **LaB1_Rod_fwd** | accctcgtaaaggtctagagaccatggactacaaagac | Cloning LaB1 ΔCaaX |
| **LaB1_CaaX_del_rev** | ccgtttaaactcattactaactagcttctattggatgctcttg | Cloning LaB1 ΔCaaX |
| **Progerin_fwd** | accctcgtaaaggtctagagaccatggactacaaagacgatgacgacaagatggaga ccccgtcccag | Cloning Progerin |
| **Progerin_rev** | ccgtttaaactcattactaattacatgatgctgcagttctgg | Cloning Progerin |
| **LaA_rod_4_chimera_ fwd** | accctcgtaaaggtctagagaccatggactacaaagacgatgacg | Cloning LaA H+R-B1 Tail |
| **LaA_rod_4_chimera_ rev** | tgctaacactgctgctgcggctctcagt | Cloning LaA H+R-B1 Tail |
| **LaB1_Tail_4_chimera _fwd** | ccgcagcagcagtgttagcatctctcattc | Cloning LaA H+R-B1 Tail |
| **LaB1_Tail_4_chimera _rev** | ccgtttaaactcattactaactacataattgcacagcttc | Cloning LaA H+R-B1 Tail |
| **LaB1_rod_4_chimera _fwd** | accctcgtaaaggtctagagaccatggactacaaagac | Cloning LaB1 H+R – A Tail |
| **LaB1_rod_4_chimera _rev** | gctgtgagaaactactcgcctctgattc | Cloning LaB1 H+R – A Tail |
| **LaA_Tail_4_chimera_ fwd** | ggcgagtagtttctcacagcacgcacgc | Cloning LaB1 H+R – A Tail |
| **LaA_Tail_4_chimera_ rev** | ccgtttaaactcattactaattacatgatgctgcagttctgg | Cloning LaB1 H+R – A Tail |
| **LaA_rod_4_chimera_ fwd** | accctcgtaaaggtctagagaccatggactacaaagacgatgacg | Cloning LaA H+R-NE81 Tail |

| | | |
|---|---|---|
| **LaA_rod_4_chimera_rev** | tgctaacactgctgctgcggctctcagt | Cloning LaA H+R-NE81 Tail |
| **NE81_Tail_fwd** | ccactgagagccgcagcagcgcagcagctaatcaggatc | Cloning LaA H+R-NE81 Tail |
| **NE81_Tail_rev** | ccgtttaaactcattactaattacatgatcagacaatttgatttc | Cloning LaA H+R-NE81 Tail |
| **LaB1_Rod_fwd** | accctcgtaaaggtctagagaccatggactacaaagac | Cloning LaB1 Head-ARod+Tail |
| **LaB1_Head_rev** | ctccttctcctggagccg | Cloning LaB1 Head-ARod+Tail |
| **LaA_Rod/Tail_fwd** | cgcggctccaggagaaggaggacctgcaggagctcaatg | Cloning LaB1 Head-ARod+Tail |
| **LaA_Rod/Tail_rev** | ccgtttaaactcattactaattacatgatgctgcagttctg | Cloning LaB1 Head-ARod+Tail |
| **LaB1_Head_fwd** | accctcgtaaaggtctagagaccatggactacaaagacgatg | Cloning LaB1Head-ARod |
| **LaB1_Head_rev** | ctccttctcctggagccg | Cloning LaB1Head-ARod |
| **LaA_Rod_4_HS_fwd** | cgcggctccaggagaaggaggacctgcaggagctcaatg | Cloning LaB1Head-ARod |
| **LaA_Rod_4_HS_rev** | ccgtttaaactcattactaattagctgctgcggctctc | Cloning LaB1Head-ARod |
| **LaA_Head_fwd** | accctcgtaaaggtctagagaccatggactacaaagacgatg | Cloning LaAHead-B1Rod+Tail |
| **LaA_Head_rev** | ctccttctcctgcagccg | Cloning LaAHead-B1Rod+Tail |
| **LaB1_Rod/Tail_fwd** | cccggctgcaggagaaggaggagctgcgcgagctcaatg | Cloning LaAHead-B1Rod+Tail |
| **LaB1_Rod/Tail_rev** | ccgtttaaactcattactaactacataattgcacagcttctattggatg | Cloning LaAHead-B1Rod+Tail |
| **LaA_Head_fwd** | accctcgtaaaggtctagagaccatggactacaaagacgatg | Cloning LaAHead-B1Rod |
| **LaA_Head_rev** | ctccttctcctgcagccg | Cloning LaAHead-B1Rod |
| **LaB1_Rod_4_HS_fwd** | ggagaaggaggagctgcgcgagctcaatg | Cloning LaAHead-B1Rod |
| **LaB1_Rod_4_HS_rev** | ccgtttaaactcattactaactaactactcgcctctgattcttcc | Cloning LaAHead-B1Rod |

**Supplemental Table 1:** List of primers used for Gibson cloning lamin expression constructs.

### Supplemental Figures

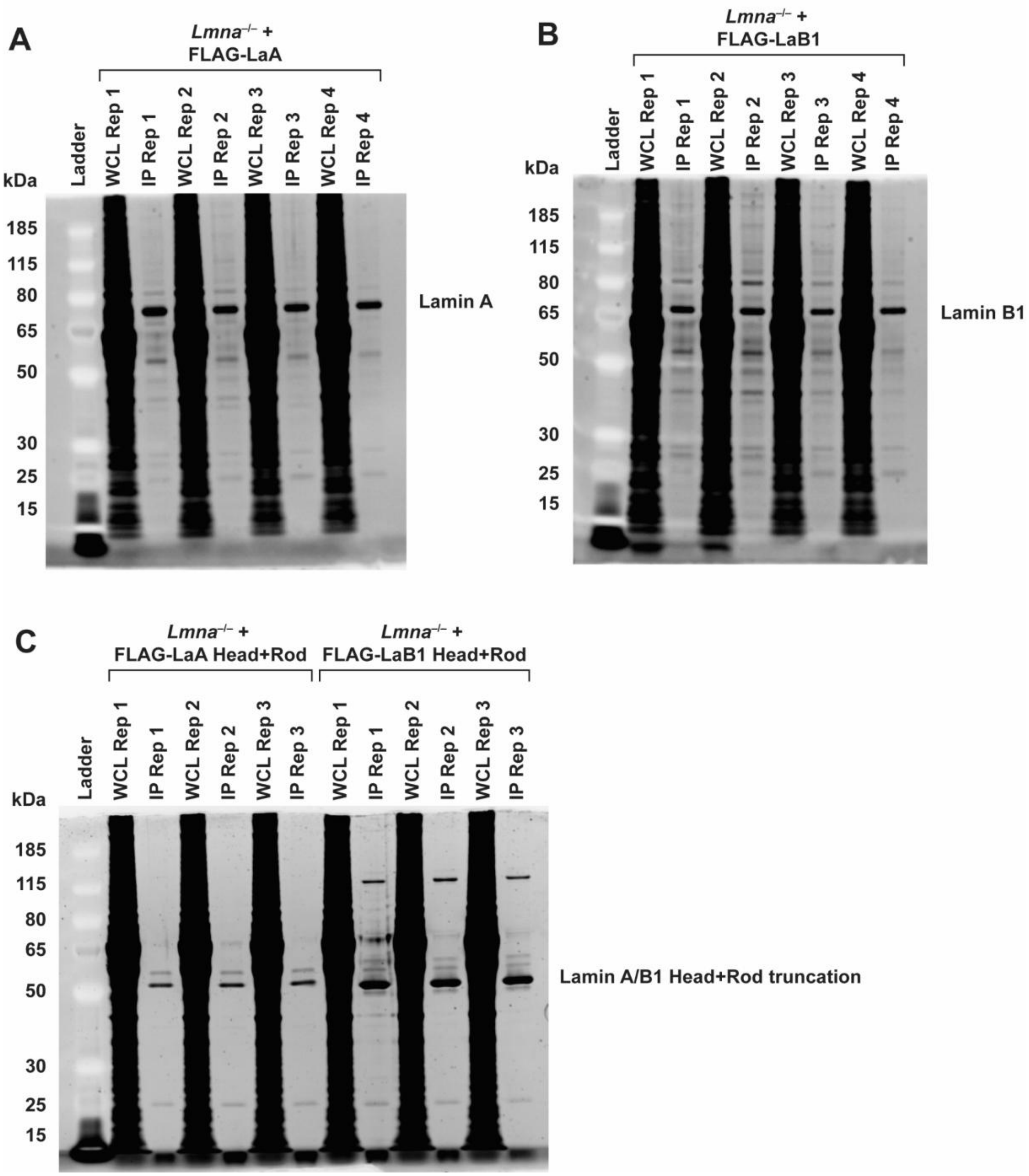


**Supplemental Figure 1: SYPRO Ruby stained gels to validate co-IP of lamin constructs.** Co-IPs were performed using anti-FLAG beads to co-IP LaA (**A**), LaB1 (**B**), or the respective Head+Rod truncations (**C**). WCL = whole cell lysate, IP = immunoprecipitation eluate. The overexposed signal in the WCL represents all proteins in the cell, and the prominent well-resolved bands in the IP conditions indicate the respective bait proteins, with fainter bands for likely interaction partners.

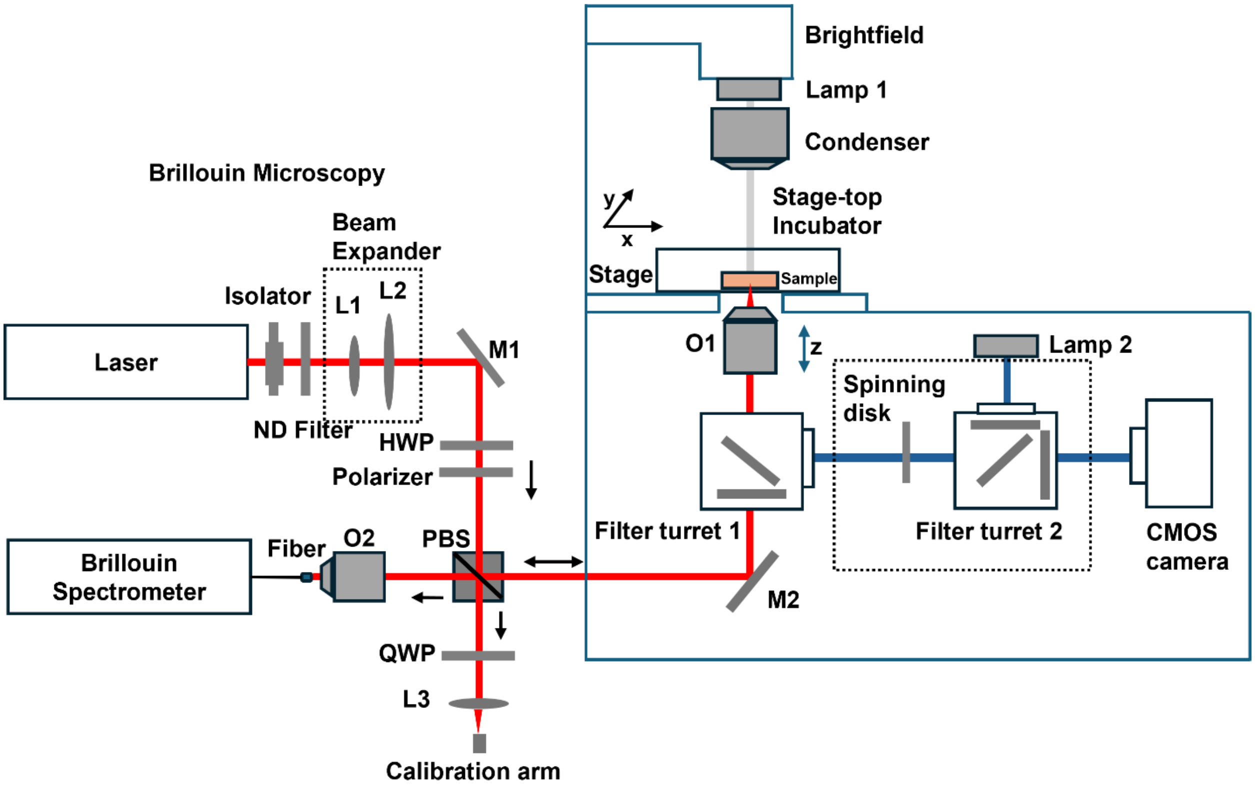


**Supplemental Figure 2: Schematic of confocal Brillouin microscopy.** HWP, half wave plate; L1, L2, L3, lens; M1, mirror; M2, dichroic mirror; ND filter, neutral density filter; O1, O2, objective lens; PBS, polarized beam splitter; QWP, quarter wave plate.
